# Improved characterization of soil organic matter by integrating FTICR-MS, liquid chromatography tandem mass spectrometry and molecular networking: a case study of root litter decay under drought conditions

**DOI:** 10.1101/2023.06.20.545455

**Authors:** Nicole DiDonato, Albert Rivas-Ubach, William Kew, Chaevien Clendinen, Noah Sokol, Jennifer E. Kyle, Carmen E. Martínez, Megan M. Foley, Nikola Tolić, Jennifer Pett-Ridge, Ljiljana Paša-Tolić

**Author notes:** Instituto Nacional de Investigación y Tecnología Agraria y Alimentaria: Madrid, Spain (Albert Rivas-Ubach).

## Abstract

Knowledge of the type of carbon contained in soils is important for predicting carbon fluxes in a warming climate, yet most soil organic matter (SOM) components are unknown. We used an integrated three-part approach to characterize SOM from decaying root-detritus microcosms subject to either drought or normal conditions. To observe broad differences in SOM compositions we employed direct infusion Fourier transform ion cyclotron resonance mass spectrometry (DI-FTICR-MS). We complemented this with liquid chromatography tandem mass spectrometry (LC-MS/MS) to identify components by library matching. Since libraries contain only a small fraction of SOM components, we also used fragment spectra cosine similarity scores to relate unknowns and library matches through molecular networks. This approach allowed us to corroborate DI-FTICR-MS molecular formulas using library matches and infer structures of unknowns from molecular networks to improve SOM annotation. We found matches to fungal metabolites, and under drought conditions, greater relative amounts of lignin-like vs condensed aromatic polyphenol formulas, and lower average nominal oxidation state of SOM carbon, suggesting reduced decomposition of carbon and/or microbes under stress. We propose this integrated approach as more comprehensive than individual analyses in parallel, with the potential to improve knowledge of the chemical composition and persistence of SOM.

**Synopsis:** Structural characterization and identifications are lacking for soil organic matter components. This study integrates molecular formula assignments and structural information from fragment ion spectra into molecular networks to better characterize unknown soil organic matter components.

**For Table of Contents Only:** 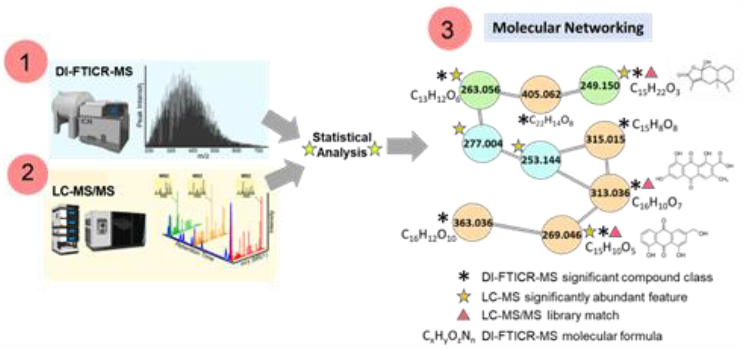

## 1. Introduction

SOM is a highly complex, heterogeneous mixture of organic components that have been modified from relatively well-known precursors, yet are mostly unidentified [1]. SOM exhibits a large range of polarities, it is not completely separable, and it is often bound to inorganic molecules and minerals, which impede characterization. SOM is an important sink in the global carbon cycle and efforts to understand what type of carbon persists in soils, and how, is of primary interest for predicting how current stocks are affected by climate change as well as how to increase the amount that can be preserved in soil over time [2]. Yet the complexity of SOM is not adequately represented in models [3-5], partly due to the lack of known SOM components of significance to carbon preservation.

Molecular formula assignment using direct infusion high-resolution mass spectrometry has become a powerful tool for untargeted analysis of natural organic matter (NOM), including SOM. The ultra-high mass resolution and mass accuracy of FTICR-MS allows for molecular formula assignment and screening via direct infusion (i.e., without the need for up-front separations). Thousands of formula assignments can be compared among samples to evaluate trends in molecular properties. Sample preparation can be minimal, data acquisition is relatively rapid (1-2 sec/acquisition), and spectra are often exceptionally rich. Although this technique can be limited by charge competition [6], instrument dynamic range[7], ambiguous molecular formula assignments[8, 9], and a lack of complete structural and chemical identification-including the unknown multiplicity of isomeric structures[10], it provides a detailed snap-shot of the diversity of SOM components in a given sample.

Alternatively, more specific molecular and structural information can be obtained by performing liquid chromatography-mass spectrometry (LC-MS), which reduces charge competition, enhances overall dynamic range, and enables molecular identification of soil components by comparing *m/z* and retention time with library databases. Liquid chromatography tandem mass spectrometry (LC-MS/MS) additionally takes advantage of precursor mass fragment ion spectra for library matching. LC-MS and LC-MS/MS platforms have been increasingly utilized to identify metabolites and lipids from environmental samples, including plant and rhizosphere extracts and, to a lesser extent, SOM [11-17]. However, only a fraction of the peaks detected can be annotated using this approach due to incomplete separation and lack of comprehensive libraries for SOM components.

When used in concert, DI-FTICR-MS and LC-MS provide an integrated approach, balancing the need for “big-picture”, compositional information on the scope of thousands of molecular formulas, with identification of a much smaller subset of components afforded by LC-MS and MS/MS. LC-MS/MS based molecular networking tools such as Global Natural Product Social Networking (GNPS) can further expand library matching and improve classification of unknowns by networking precursor masses based on the similarity of their fragmentation spectra [18-21]. Networks are especially informative for inferring structures of unknowns networked to library matches.

The use of FTICR-MS and LC-MS/MS for investigation of complex environmental samples has most often been applied to dissolved organic matter (DOM). Several studies have reported improved detection of DOM using chromatography [22, 23], quadrupolar detection [24] and increasing magnetic field strength [7, 25]. Others have reported on the extreme isomeric complexity of DOM [10, 26], and the ability to match LC-MS/MS spectra with library spectra in order to identify DOM components [27-29] and create molecular networks [30, 31]. Still only a handful of studies [32, 33] have utilized these untargeted approaches to identify metabolites from soils, likely due to additional challenges separating poorly ionizable material present in greater proportions in terrestrial samples, and the lack of appropriate libraries [6]. Only one other study to our knowledge has employed fragment ion spectra-based molecular networking for soil extracts [34]. We are unaware of other studies that have evaluated DI-FTICR-MS molecular formula assignments of soil extracts using LC-MS and LC-MS/MS based identifications and molecular networks.

In this work, DI-FTICR-MS, LC-MS, and LC-MS/MS classical molecular networking are utilized for chemical characterization of SOM formed from root detritus of a common annual grass species, *Avena barbata*, under simulated normal moisture (∼16% soil moisture, ‘control’) and drought conditions (∼8% soil moisture). Our overarching goal was to combine the complimentary information from each of these techniques for a more robust approach for SOM analysis. Library matches and networked, precursor ion masses were compared with masses detected using DI-FTICR-MS to corroborate molecular formula assignments and provide additional structural information for unknown SOM analytes. We also assessed the fraction of formulas observed with direct infusion that can be separated using LC-MS, matched with existing libraries, or networked to library matches. We focus primarily on this novel approach (Figure 1), to illustrate the potential to improve the number of identifications and knowledge of SOM constituents that may be significant for SOM preservation in soils.

**Figure 1:**
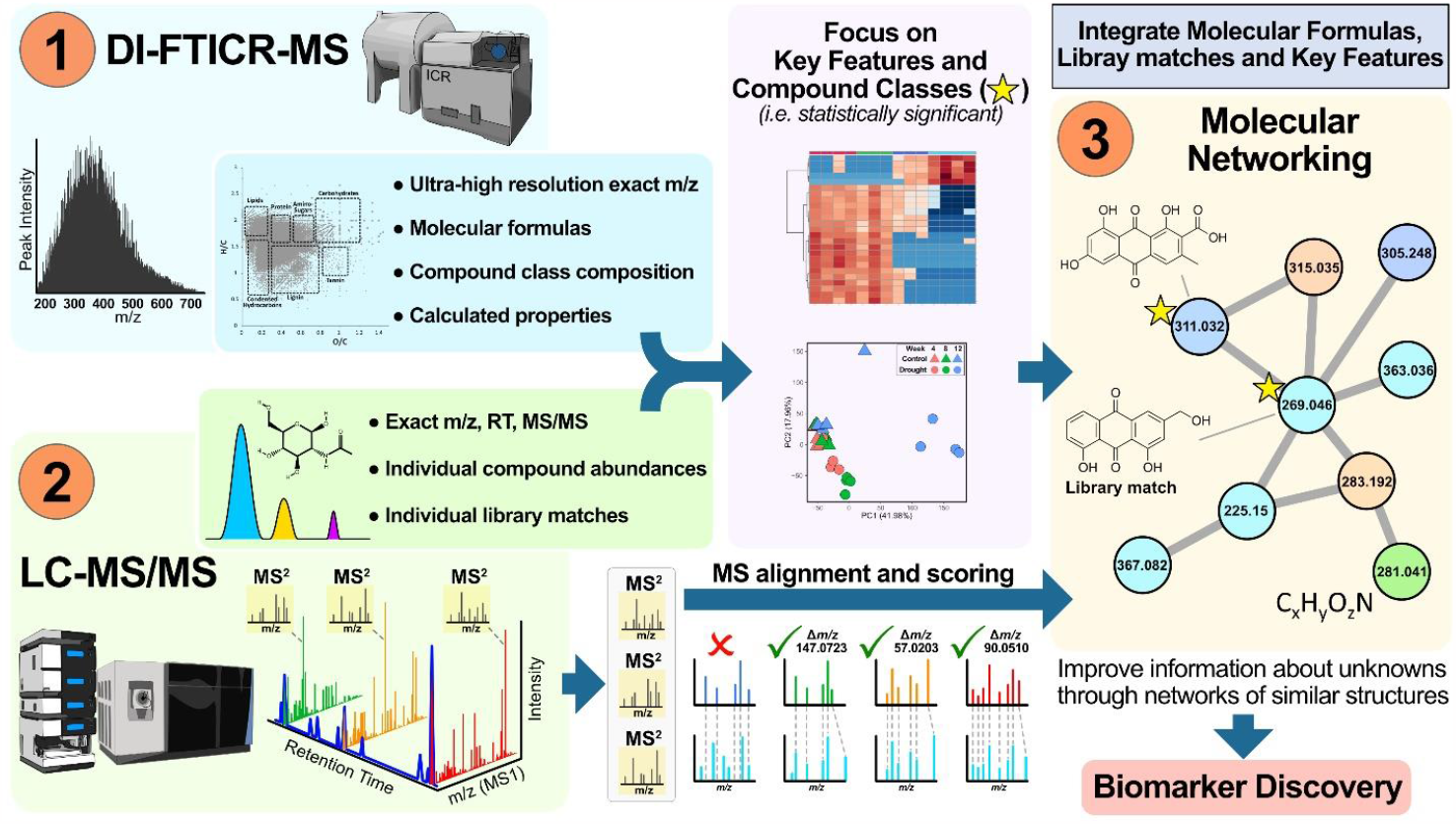
Integration of SOM measurements using 1) molecular formulas detected with DI-FTICR-MS, 2) library matches identified with LC-MS/MS and 3) molecular networks based on MS/MS spectra of unknown molecular formulas and library matches. Statistical analysis of both DI-FTICRMS and LC-MS/MS features focuses our analysis on SOM components with significantly different abundances between treatments or significantly different molecular class composition, creating a more targeted approach to SOM characterization by integrating individual non-targeted techniques.

## 2. Materials and Methods

### 2.1 Soil samples

Details of the soil collection site and growth chamber conditions are described in the SI. Briefly, soil was collected from the ‘A’ mineral horizon at the University of California Hopland Research and Extension Center in Hopland, CA and approximately 65 g of soil was mixed with *A. barbata* root litter fragments (1-5 mm) and packed into acrylic soil microcosms (56 x 56 x 76 linear cm) that were placed in greenhouse growth chambers maintained at one of two moisture treatments: ‘normal moisture’ (∼16% ± 0.3 gravimetric soil moisture; mean ± standard error) or ‘drought’ conditions (∼8% ± 0.5 gravimetric soil moisture). Microcosms were destructively harvested from each treatment (n = 4) to collect soil samples at 4, 8, and 12 weeks.

Soil samples were dried, ground, weighed, and analyzed for % total C on an elemental analyzer (EA-IRMS; Costech ECS 4010, Costech Analytical Technologies, Valencia, CA, USA) at the Yale Analytical and Stable Isotope Center (New Haven, CT, USA). Across time points, the average percent carbon values for the drought treatment (2.31 +/-0.12%) and control (2.35 +/-0.39) samples were not statistically different (Table S1).

### 2.2 Extractions

Soil samples were weighed, flash frozen, freeze-dried and shipped to the Environmental Molecular Science Laboratory (EMSL, Richland, WA) for extraction and analysis. 1) A modified MPLEx extraction protocol was used to extract SOM from 1g of dry soil per sample and is detailed as supporting information [35, 36]. This produced 3 extracts for each sample: water, methanol and chloroform, for greater coverage of SOM components. 2) Salts and impurities that could interfere with MS analysis were removed from the water extract using solid phase extraction (SPE) and SOM was eluted in methanol [37]. Water-extracted SOM eluted in methanol from step (2) and methanol and chloroform extracts from step (1) were analyzed individually with DI-FTICR-MS. The same extracts were combined for LC-MS/MS analysis. Details of the extraction procedures can be found in the SI.

### 2.3 Direct Infusion ESI-FTICR-MS

A 12 Tesla Bruker SolariX FTICR-MS housed at EMSL was used for analysis of water, methanol and chloroform extracts separately in negative ion mode. Details of the method are provided as supporting information. Formulas were assigned using Formularity software [38] with N<=2, S=0 and P=0, initially for high confidence assignments (average error < 0.2 ppm). Molecular class distributions were based on percentage of formulas with elemental ratios within the H/C and O/C ranges for major compound classes [38, 39]. Differences in calculated mean NOSC values and mean class distributions were evaluated for statistical significance using the *Anova* function as part of the “ftmsRanalysis” package in R developed at PNNL [40].

Classifications provide a useful comparison of formula types based solely on O/C and H/C ratios; the presence of N, S or P was not considered for compound classifications. For example, molecular formulas classified as “protein-like” will have similar O/C and H/C ratios as proteinaceous compounds (i.e. amino acids, peptides, etc.), but may or may not contain nitrogen. Inferences to the origin of compound classes require additional information. Though the classes are putative, we drop the “-like” designation for clarity in figures.

*m/z* and assigned formulas from each extract were combined into a composite list of components by sample. For comparison between peaks detected with DI-FTICR-MS and LC-MS, all *m/z* peaks (z=1) were rounded to *m/z* 0.001 (1 mDa). N, S and P restrictions on direct infusion molecular formula assignments were relaxed for comparison with LC-MS/MS library matches containing N, S and P.

### 2.4 LC-MS and MS/MS analysis of soil extracts

Metabolomics and lipidomics platforms at PNNL with libraries containing plant and microbial metabolites and lipids were used for this study. Water, methanol and chloroform extracts were combined prior to metabolite analysis as described in the SI. Combined extracts were analyzed with a Thermo Vanquish High Pressure Liquid Chromatography (HPLC) (Thermo Fisher Scientific, Waltham, Massachusetts, USA). SOM components were separated by injecting 5 µL of each sample onto a Hypersil gold C18 reversed-phase column (150×2.1 mm, 3 μm particle size; Thermo Scientific, Waltham, Massachusetts, USA) operating at 30°C. LC mobile phases consisted of 0.1% formic acid in water (A) and 0.1% formic acid in acetonitrile/water (90/10) (B). A 35-minute gradient elution was performed and is detailed in the SI. High-resolution mass spectrometry (HRMS) was performed on a LTQ Orbitrap Velos equipped with a heated electrospray ionization (HESI) source (Thermo Fisher Scientific, Waltham, Massachusetts, USA). Details of HRMS measurements are provided in the SI.

Negative and positive LC-MS RAW files of metabolite data were processed with MZmine version 2.51 [41]. Data processing details are described in the SI. Datasets from negative and positive ionization modes were combined into one for statistical analyses. All LC-MS metabolite statistical analyses were performed in R version 4.1.1 (R Core Team 2020). One-way ANOVAs were calculated with the *aov* function, from “STATS” package (R Core Team 2020) to contrast drought versus control soil samples at the final time point (week 12) for each individual detected feature. Principal component analysis (PCA) was performed on the entire metabolomic fingerprints of soils including both categorical factors (moisture treatment and time point) with functions *imputePCA* and *PCA* from “missMDA” [42] and “FactoMineR” [43] packages, respectively.

Chloroform-only extracts were analyzed for lipids as outlined by Kyle et al. and details for this analysis can be found in the SI [44].

### 2.5 SOM Library Matching

We used two libraries for metabolite identifications: our in-house metabolomics library and external GNPS libraries. Metabolite masses with unique retention times (RT) were matched according to the exact monoisotopic mass and RT of standard compounds in our in-house library which contains over 600 standards of common plant and microbial metabolites. According to the metabolomics standards initiative this compound match corresponds to a second level identification [45]. Fragmentation spectra of precursor ions were used to validate identifications but were not required for compound matching. Lipids were identified using LIQUID as outline in Kyle et al [44]. Briefly, confident identifications were assigned using accurate mass, retention time and fragment ions. The library in LIQUID contains over 45,00 lipids across 80+ types of lipids from bacteria, plants, fungi and mammals.

Community based external Global Natural Product Social Networking (GNPS) spectral libraries were compared with experimental fragment ion spectra for precursor masses selected via DDA in both metabolomics and lipidomics LC-MS/MS datasets. Library matching was performed separately for datasets from each platform and ionization mode as part of the classical molecular networking workflow offered by GNPS.

### 2.6 GNPS Classical Molecular Networking

Raw files from both metabolomics and lipidomics LC-MS/MS analyses were converted to mzML and uploaded to the GNPS portal for fragment spectra clustering, comparison with libraries and networking. Parameters used for GNPS classical networking can be found in Table S3.

## 3. Results and Discussion

### 3.1 Direct infusion FTICR-MS characterizes broad changes in SOM composition

SOM extracts were assessed for the relative differences in average chemical properties and average abundance of compound classes calculated from formula assignments. Average formula assignments for the detritusphere SOM are presented in Table S4, along with calculated properties, including mean nominal oxidation of carbon (NOSC). NOSC is a measure of how much energy is required to oxidize a molecule to CO_2_, ranging from -4 (CH_4_) to +4 (CO_2_), with lower values requiring more energy than higher values. Mean NOSC values varied primarily with polarity of the extractant (water>methanol>chloroform) as expected (Table S4); molecules extracted with more polar solvents contain more heteroatoms (which impart polarity) and were found to be more oxidized, having higher NOSC values. At 4 and 8 weeks, no significant difference in NOSC was detected between drought and control treatments for these low carbon (∼2%), high mineral content samples. However, mean NOSC values for the water extractable carbon in the drought treatment were found to decrease relative to the control over the course of the experiment and this difference was significant by 12 weeks (P < 0.01, Figure 2a). In general, lower NOSC values indicate that more energy is required to convert compounds to CO_2_ in the drought compared to the control by the end of the experiment. Consistent with decreased NOSC values, mean H/C ratios increased, and mean O/C ratios decreased in the drought relative to the control and this difference was also significant at the last time point (Week 12 H/C, P< 0.01, Week 12 O/C, P<0.05).

**Figure 2.**
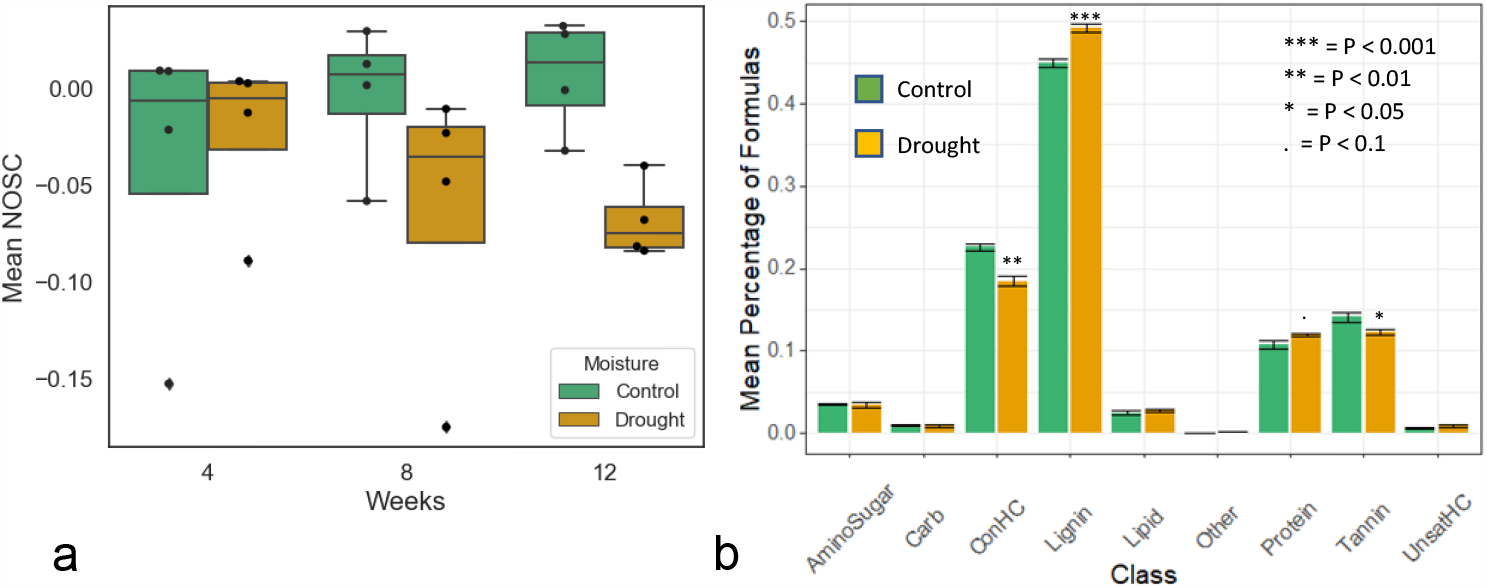
a) Mean NOSC values for molecular formulas detected in water-soluble carbon extracted from detritusphere samples at week 4, 8 and 12 (n = 4) b) Class distributions of water extractable carbon in detritusphere at week 12 (n = 4). Asterisks in b) indicate significance levels for differences between means determined via *Anova* (*Carb = Carbohydrate, ConHC =Condensed hydrocarbon, UnsatHC = Unsaturated hydrocarbon)*

By the end of the experiment (week 12), the percentage of molecular formulas plotting in the tannin-like, protein-like, condensed hydrocarbon and lignin-like regions in the drought treatment were statistically different from the control (Figure 2b). The most significant differences in molecular class composition were the greater proportion of lignin-like (p<0.001) and the smaller proportion of condensed hydrocarbon-like compounds (p<0.01) found in the drought treatment after 12 weeks (Figure 2b and S1). Though classes of compounds are based on elemental ratios alone and are not exclusive of the types of compounds that could be present [46], our interpretation is that the overall SOM composition changed, and compound classes that change significantly provide a focus for investigating changes occurring at the molecular level due to drought. For example, the shift in molecular formula composition could represent a change in the decomposition of organic matter under drought conditions. Compounds with lower NOSC values may be left behind by microbes in the drought treated soils that do not have, or cannot allocate, the enzymes necessary for their decomposition [47, 48]. Alternatively, this may also suggest a change in metabolite production as a result of drought induced stress or a shift in microbial community composition [49]. To better understand the observed trends, we used LC-MS for metabolite identification to determine if components could be separated and identified, particularly those compound classes with the most significant difference between treatments (lignin-like, condensed hydrocarbon, etc.). Since drought had the most pronounced effect on SOM at week 12, we focus primarily on this time point to investigate differences between moisture treatments, and we concentrate on negative ion mode analytes to highlight how library identifications corroborate and compliment direct infusion formula assignments.

### 3.2 Library Matches Validate DI-FTICR-MS Formulas

Our in-house library was used to identify components of detritusphere SOM based on retention time and accurate mass measurements of standards common to plant and microbial metabolites. For comparison with DI-FTICR-MS, primarily performed in negative ion mode, 1269 compounds, or “features” with unique mass and retention time, ranging from *m/z* 74-619, were detected in detritusphere samples from negative ion mode LC-MS spectra alone. This led to 42 matches to the in-house library (Table S5a and c). GNPS community-based libraries were also used to identify SOM components by comparing experimental fragment ion spectra with library spectra. 739 precursor ion fragmentation spectra were collected from negative mode spectra, resulting in 36 putative matches to GNPS libraries for detritusphere samples at the end of the 12-week experiment. Although non-targeted LC-MS studies commonly match only a small number of compounds using metabolite libraries [46], the relatively small number of matches found in the current study also directly reflects the vast complexity of soil organic matter and the fact that libraries contain predominantly primary metabolites. Only 7 compounds were matched by both our in-house and GNPS libraries, hence the total number of library matches almost doubled by searching GNPS libraries, in addition to the in-house library alone.

As with the FTICR-MS results presented above, the most significant difference between metabolites in the drought and control samples was found at the end of the experiment (week 12) (Figure 3a) where drought soil samples were most separated along PC1 from control samples and earlier time points. Though statistical analysis of LC-MS data is based on metabolite relative abundance from both ion modes and FTICR-MS data considered only the presence or absence of a peak from negative ion mode, both methods resulted in similar trends. We also found a greater percentage of LC-MS compounds (27.6% vs 10.8%) with significantly higher abundance in the drought treatment, compared to the control, at the end of the experiment (Figure 3b). This could suggest decreased microbial assimilation of SOC under drought conditions, and/or it could indicate the production of drought-induced metabolites.

**Figure 3.**
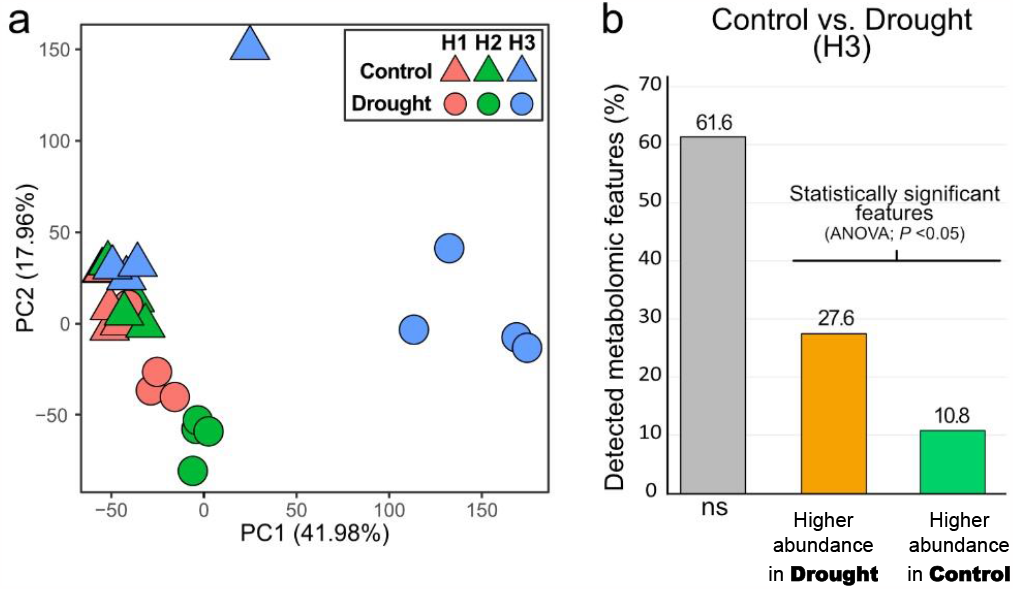
a) Principal component (PC)1 vs PC2 for the principal component analysis (PCA) case plot of the LC-MS metabolite fingerprint from detritusphere samples under control (triangles) and drought conditions (circles) at week 4 (red), 8 (green) and 12 (blue) from both positive and negative mode spectra (**a**). Percentage of metabolic features non-significantly changing (*ns*) and significantly changing (*P* <0.05) with higher abundance in either soils under control or drought conditions (**b**).

Library matches with significantly different abundances between treatments can inform the observed trends, yet since matches are few, we used multiple libraries as well as DI-FTICRMS molecular formulas to improve coverage and to cross-check our findings. Putative matches found to have significantly different abundances between moisture treatments (Tables S5a-c) include 14 in-house library matches (8 from negative mode spectra including emodin, purine, leucine, adenine, L-glutamic acid, hydroxymethyl glutarate, hexoses, and adenosine) and 10 GNPS matches (including 5 matches to negative mode spectra for esculetin, an isomer of herticin A, deoxyguanosine, aloe-emodin and a phenolic acid glycoside). Significantly higher abundance of putative matches adenosine, alanine and hydroxymethyl glutarate under drought may be related to microbial regulation of energy metabolism while others, such as polyphenolic compounds, as discussed in more detail in Section 3.4, may be related to microbial defense [49, 50]. In-house matches that had significantly higher abundance in the control (hypoxanthine, adenine, guanine, and rac-glycerol 1-myristate) (Table S5a), could be related to greater microbial activity and carbon assimilation under normal conditions [51, 52].

A large portion of compounds with significantly higher abundance under drought conditions remain unknown. Yet approximately 10% of those detected in negative ion mode were also detected (+/-1 mDa) with DI-FTICR-MS and assigned molecular formulas using restricted criteria (N<=2, S = 0, P= 0). These plot primarily with lignin-like and lipid-like H/C and O/C ratios (Figure S2).

The remaining unknown LC-MS features for which molecular formulas could be assigned (+/-1 mDa) using DI-FTICRMS are also plotted in Figure 4. Included are 64% (231) of the 360 unique precursor-masses with acquired fragmentation spectra that were also detected (+/-1 mDa) using DI-FTICR-MS (about half of the precursor masses, see Figure S3). These precursor masses cover a range of polarities and molecular size, as evidenced by their retention times (Figure S3).

**Figure 4.**
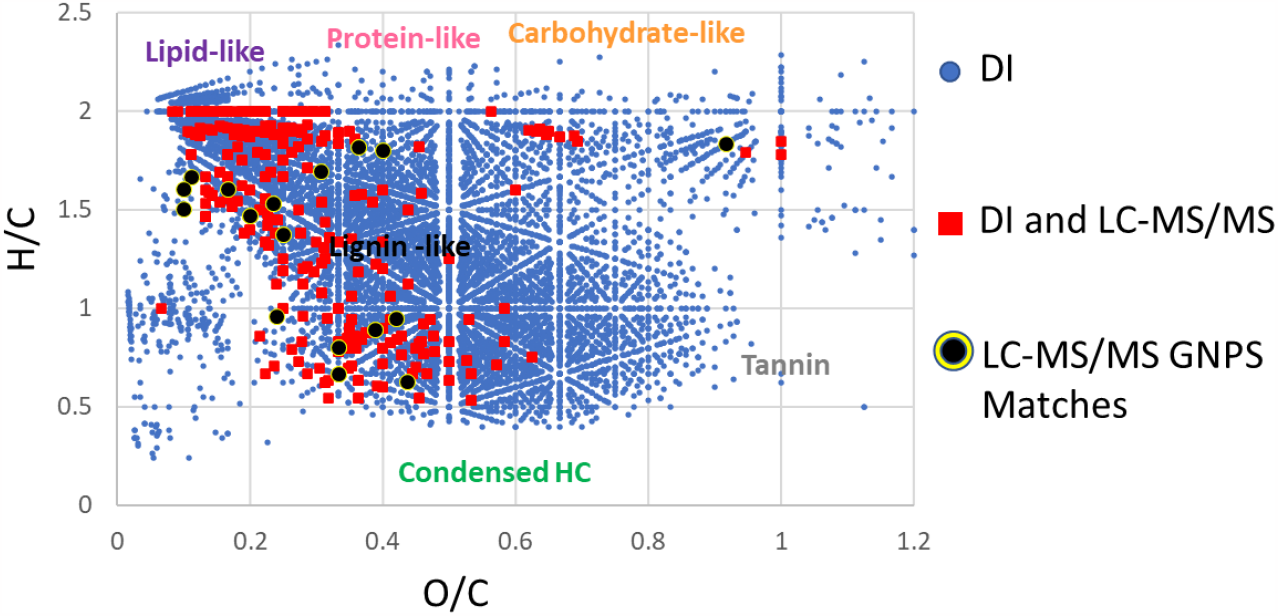
van Krevelen diagram of DI-FTICR-MS molecular formulas assigned from SOM extracts at week 12 (drought + control) (blue). Molecular formulas for precursor masses that were also detected with LC-MS (+/-1 mDa) and acquired MS/MS spectra are overlayed in red. Features with fragmentation spectra that match with GNPS library spectra are highlighted in black and tabulated in Table S5c.

Of 42 putatively identified in-house library matches, only 9 were within the *m/z* range detected with DI-FTICMRS (200-900 *m/z*: thymidine, cytidine, palmitoleic acid, adenosine, inosine, emodin, disaccharides, trisaccharides and usnic acid). Yet, all 9 compounds were also detected within 1 mDa in at least one sample extract analyzed with DI-FTICR-MS and all were assigned molecular formulas that agreed with the library match (+/-0.23 ppm) (Table S5c). A slight bias of the in-house library towards low mass metabolites (58% with *m/z* < 200) likely contributes to the low overlap between in-house library matches and DI-FTICR-MS measured masses.

There were a greater number of GNPS matches with *m/z* >200 and detected with DI-FTICR-MS within 1 mDa of the LC-MS *m/z* (26 of the 36 GNPS matches). 17 of these were assigned molecular formulas using strict formula assignment criteria (N<=2, S=0, P =0) (Table S5c, Figure 4 and S3). All were assigned the same molecular formula as the GNPS match, with a maximum mass error of 0.23 ppm (Table S5c), thus validating molecular formula assignments. Matches span the regions of the van Krevelen diagram, indicating lipid-like, condensed hydrocarbon, and lignin-like classes of compounds. There were no conflicts between DI-FTICR-MS molecular formulas and library matches.

Library matches with lignin-like elemental ratios may provide some indication of the types of compounds, as a class, found to be present in greater proportion in the drought treatment using DI-FTICR-MS. GNPS matches with elemental ratios similar to lignin-like compounds include phenolics such as usnic acid, gallic acid, syringic acid, atronorin, animicin a, an isomer of the fungal metabolite isopestacin [53] (3-(2,6-dihydroxyphenyl)-4-hydroxy-6-methyl-3H-2-benzofuran-1-one) and similar compounds such as juarezic acid and an isomer of the sesquiterpene medicinal plant metabolite herticin A [54] ((4aR,5S,8aS,9aR)-9a-hydroxy-3,4a,5-trimethyl-5,6,7,8,8a,9-hexahydro-4H-benzo[f][1]benzofuran-2-one) (Table S5c). In-house library matches with H/C and O/C elemental ratios for this molecular class also include usnic acid, inosine, adenosine, cytidine and thymidine. Based on LC-MS, adenosine was found to have a significantly higher abundance (P<0.05) in the detritusphere soils under drought, and usnic acid, inosine and cytidine were also detected at higher abundance (P<0.1).

#### 3.2.1 Quality and validation of GNPS matches

Cosine similarity scores and the number of matching fragment ion peaks are two GNPS measures to assess the quality of a library match. Cosine similarity scores are a mathematical measure of similarity based on fragment ion *m/z* and intensity and precursor *m/z*, where a score of 0 represents no similarity and 1 denotes identical fragmentation spectra. Scores for GNPS matches in this study are listed in Table S5b and ranged from the default recommended minimum score of 0.7, to 0.97.

There is a tradeoff between the threshold for the minimum number of matching fragment ions and the number of matches obtained. Plausible matches with few matching fragment ions, such as esculetin, endocrocin, and aloe emodin (cosine scores 0.92, 0.84 and 0.79) would be excluded by using a stringent threshold of 6 minimum matching fragment ions. Alternatively relaxing this threshold can lead to more questionable matches that require validation. Lowering the minimum matching fragment ion threshold from 6 to 3, generated 26 additional putative matches (up from just 10), containing 3 to 11 matching fragment ion peaks. We were able to independently substantiate at least one match (aloe-emodin, cosine score 0.79) with as little as 3 matching fragment ion peaks using our in-house library authentic standard. The remaining 6 of the 7 GNPS matches verified by our in-house library were matched to GNPS library spectra with at least 4 and up to 7 matching fragment ion peaks (Table S5d). In the absence of standards, manual inspection is required and often inconclusive. However, integrating information from molecular networks and accurate mass molecular formulas can help evaluate a match, as described in the next section.

### 3.3 Molecular networks provide structural information for unidentified DI-FTICR-MS formulas

Molecular networks allow us to group unknown precursor masses with similar fragmentation patterns together. This is built on the concept that structurally related molecules exhibit similar fragmentation under similar conditions [18, 19]. Fragment ion spectra of molecules with similar building blocks can have several shared fragment ion peaks corresponding to the structural entities they hold in common. Alternatively, fragment ion peaks with similar ion intensity distributions may be shifted by a common delta mass representing the piece of the molecule that is different. A combination of these scenarios could also occur.

The complete molecular network for metabolites detected at week 12 from drought and control soil treatments is shown in Figure 5a. Since DI-FTICR-MS measurements indicated that lignin-like and condensed hydrocarbon compounds may be of interest to this study (Figure 2b), we zoom in on a molecular subnetwork where several masses detected with DI-FTICR-MS were matched to GNPS library compounds with polyphenolic structures (Figure 5, Table S6). Figure 5d shows precursor masses *m/z* 269.0455 and *m/z* 313.0354 that were matched to GNPS library spectra for aloe-emodin and endocrocin, respectively and share a similar intensity distribution for 5 fragment ion peaks (cosine score of 0.95) each shifted by 43.9898 Da, the mass of a CO_2_ group. Their DI-FTICR-MS assigned molecular formulas, C_15_H_10_O_5_ (−0.5 mDa) and C_16_H_10_O_7_ (+0.4 mDa), respectively, and their molecular structures corroborate this difference. They share the same anthranoid scaffold, endocrocin differing from aloe-emodin by a carboxylic acid group at position 3, a methyl group in place of a hydroxymethy group at position 2 and a hydroxyl group at position 7 [55]. As mentioned above, the GNPS match to aloe-emodin was also substantiated by the in-house library authentic standard emodin, thus increasing our confidence in this and the putative match to endocrocin, a secondary fungal metabolite [56].

**Figure 5.**
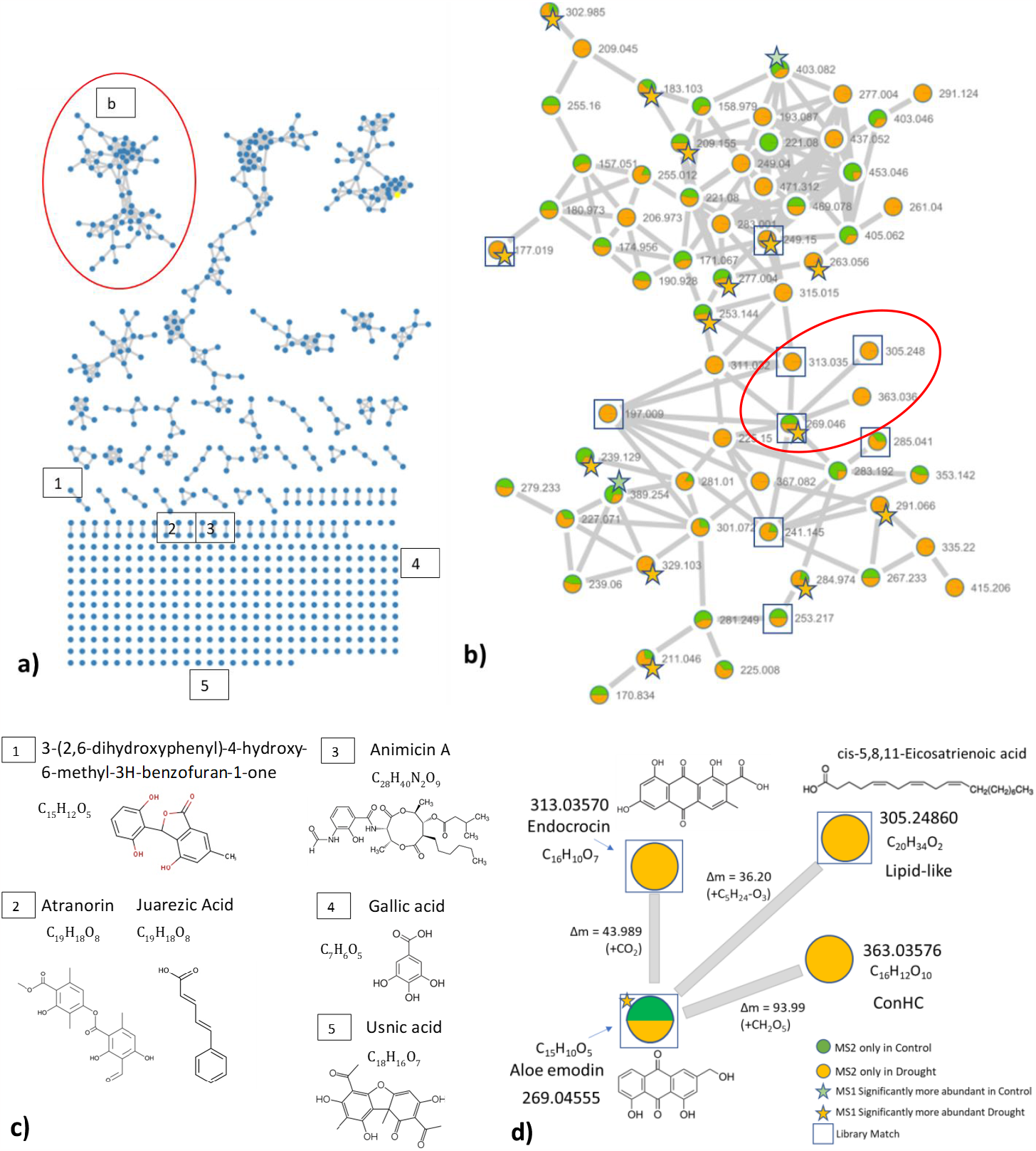
Information gathered from SOM molecular networks is improved by mapping significantly abundant precursor masses (P<0.05, determined via ANOVA) and molecular formulas for precursor masses also detected (+/-1mDa) with DI-FTICR-MS. a) Complete molecular network of nodes representing precursor masses detected in drought and control treatments of detritusphere soil samples at week 12. Masses with fragmentation spectra similarity cosine score > 0.7 are connected by gray edges. b) Subnetwork from a) containing GNPS matches to polyphenolic compounds and molecular formulas for precursor masses also detected with DI-FTICRMS. c) Several examples of matches to GNPS libraries that have lignin-like elemental ratios d) Cluster of masses with similar fragmentation spectra to aloe-emodin and assigned molecular formulas using DI-FTICR-MS.

3 of the 4 precursor masses in Figure 5d were found to be structurally similar to phenolic compounds based on LC-MS/MS matching and DI-FTICRMS formula assignments. One exception, *m/z* 305.2486 matched to an aliphatic compound with a lipid-like molecular formula. One possible explanation is that either the match or the network connection is incorrect.

Alternatively, the 5 fragment ions (ranging from 97-247 *m/z*) shared between the experimental spectrum and the library match could represent a piece of the molecule that has been fragmented and rearranged under CID conditions to produce fragments with different structures from the parent ion (i.e. reversible Diels Alder Reaction).

To illustrate how molecular networking can further complement DI-FTICR-MS molecular formula assignments, even when the number of GNPS matches are few, we examine a third precursor mass in this region, *m/z* 363.0358, that was independently detected with DI-FTICR-MS, assigned the molecular formula C_16_H_12_O_10_, but was not matched to either library. It’s H/C and O/C elemental ratios fall in the condensed hydrocarbon-like class of compounds. Without molecular networking, this would be the extent of information we have about this compound. However, this subnetwork shows that it also shares with aloe-emodin a similar fragment ion intensity distribution of 6 peaks, 2 with the same *m/z* and 4 shifted by 93.9902 Da, or CH_2_O_5_ from aloe-emodin. This agrees with the DI-FTICR-MS molecular formula assigned and provides additional confidence that this molecular formula was assigned correctly. Further, we learn that the structure of this molecule is likely similar to that of aloe-emodin, with an additional carbon and oxygen-containing groups. Another precursor mass *m/z* 291.0663 was found to have significantly higher abundance under drought conditions (Table S6) but was not matched to either library. Using this network and the DI-FTICRMS assigned molecular formula for this *m/z*, a condensed hydrocarbon-like compound C_18_H_12_O_4_, we find that it has a similar fragmentation spectrum to the dicarboxylic acid Z-2-octylpent-2-enedioic acid (*m/z* 241.1445) and differs from it by + C_5_ – H_10_ (*m/z* 49.9208), possibly indicative of a ringed structure with carboxylic acid groups.

The entire subnetwork contains a total of 64 precursor masses; 68% were detected within 1 mDa using DI-FTICR-MS and 52% were assigned molecular formulas using stringent criteria (N<=2 and S, P = 0) (Table S6). Several precursor masses were assigned molecular formulas with lignin-like or condensed hydrocarbon -like elemental ratios. Many of these shared similar fragmentation spectra (i.e. connected by edges with similarity score >0.7) to one of the 8 GNPS library matches. 6 GNPS library matches in this subnetwork with *m/z* > 200 were also detected with DI FTICR-MS (+/-1 mDa) and assigned molecular formulas in agreement with the library match. The number of shared fragment ions, library precursor mass error, and score for each match is noted in Table S6. Matches include formulas classified as lignin-like, condensed hydrocarbon, protein and lipid-like. 8 additional unknowns that had significantly higher abundance under drought conditions (Table S6) were also assigned molecular formulas based on DI-FTICR-MS measurements, similarly classified into these molecular classes of compounds and their orientation in the network is shown in Figure 5b. By comparing untargeted data-dependent LC-MS/MS features with DI-FTICR-MS, we found 3-30% of direct infusion-assigned molecular formulas per sample could be networked using GNPS classical molecular networking.

### 3.4 Biological Relevance of Library Matches and Molecular Formulas

The limited number of library matches provide an incomplete picture of SOM composition, however together with findings of overall changes in molecular class composition and reduced average NOSC with drought, we can discuss some possible biological processes underlying our observations. Microbial analyses from this same study indicated that microbial exoenzyme activity was reduced under drought, as was cumulative ^13^C carbon assimilation, and fungi were the dominant community members present under both conditions [57]. It is possible that secondary fungal metabolites with condensed-ring structures such as melanins, aromatic polyketides and hydroxy anthraquinones could represent some of the condensed hydrocarbon-like (H/C < 0.8) compounds found in greater relative proportion under normal conditions [58-60]. Under drought however, decreased activity of exoenzymes necessary to decompose complex root detritus may explain the lower average NOSC, lower O/C, and higher H/C and greater proportion of lignin-like compounds [47, 48].

Alternatively, higher H/C and lower O/C compounds with lower NOSC may be produced by soil microbes as part of a stress response or redistribution of community composition under adverse conditions [58]. Top carbon assimilators in both fungal and bacterial communities were found to shift with soil moisture treatment [57]. While the amount of ^13^C labeled mineral associated organic matter (MAOM) from active microbes was similar under drought and control conditions after 12 weeks, the composition differed [57], suggesting that production and fate of different SOM components may impact the type of carbon that persists in these soils under drought.

Library matches support the presence of polyphenols with both condensed aromatic and lignin-like elemental ratios, including fungal and lichen metabolites, that may be important under drought conditions. A few of these matches with significantly higher abundance under drought conditions include in-house library matches to rhapontin and emodin, as well as GNPS library matches to aloe-emodin, esculetin, an isomer of herticin A and an unidentified phenolic acid glycoside (P<0.05, Figure 5, Table S5c). Emodin is a polycyclic aromatic anthraquinone produced by soil fungi that grow on decaying organic matter [61, 62] and phenolic acid glyocides are thought to be a reserve pool for additional bioactive compounds [63]. Rhapontin, esculetin and herticin A also confer antimicrobial and antioxidant properties and are commonly found in medicinal plants [54, 64, 65]. On the other hand, significantly higher abundance of nucleic acids adenine and guanine in the control could be related to higher microbial activity under normal moisture conditions [51, 52, 57].

Other library matches, such as syringic and gallic acid, have lignin-like elemental ratios and are known degradation products of lignin [66]. Syringic acid is also a metabolite linked to antioxidant properties of drought tolerant plants [67]. Usnic acid and atronorin are two of the most common secondary metabolites of fungal-cyanobacteria symbiotic lichen species, almost exclusively found in lichen which are known to be resistant to drought, and rarely found in mosses and other higher plants [68, 69]. Another GNPS match (*m/z* 271.0612) with lignin-like elemental ratios is an isomer of the fungal metabolite isopestacin [53]. Animicin A is a benzoic acid derivative and secondary metabolite of streptomyces saprophytic soil bacteria that has use as a fungicide, insecticide and miticide [70]. *m/z* 407.0529, detected in the soil chloroform extracts analyzed for lipids matched to another lichen-specific secondary metabolite, chloratranorin, also with lignin-like elemental ratios.

An additional 100 lipids from negative ion mode spectra and 202 lipids from positive mode spectra provide further evidence for potential community members present in the detritus soils, which, as described in detail in the SI, supports the metabolite findings. Statistically significant library matches from both metabolomic and lipidomic workflows and molecular formula assignments from DI-FTICR-MS all suggest fungal and lichen metabolites, could be a factor to distinguish organic matter decay under drought conditions from control conditions. Lichen and fungi are resilient organisms when exposed to drought and are often one of the first species able to grow under harsh conditions, colonizing rocks and other sites where nutrients are less available [71]. These microbes may also be important for soil carbon preservation, as reported in a related study in which stable isotope probing was used to track carbon from ^13^C labeled decaying fungal necromass onto minerals [72]. Scanning electron microscope (SEM) and secondary ion monitoring (nanoSIMs) imaging showed most of the labeled carbon transferred to minerals originated from decomposing fungi as opposed to DOM.

Future work to characterize metabolite production by model fungal and microbial communities under drought conditions could help validate some of the observed trends. Combining our results with microbial and viral metagenomics as well as a complete carbon budget, we hope to gain more insight into the processes which drive carbon production, decomposition, and preservation under drought conditions.

### 3.5 Improved Coverage of Microbial Signatures from Integrated Analysis

In the current study, we show how LC-MS/MS based molecular networks and DI-FTICR-MS molecular formula assignments, when integrated, provide complimentary information for untargeted analysis of SOM (Figure 1). Statistical analysis of SOM components measured by DI-FTICR-MS reveals significant changes in molecular class composition and average chemical properties for a majority of compounds that cannot be identified using LC-MS/MS, whereas LC-MS normalized peak intensities highlight individual compounds with significantly different abundances that cannot be quantified using DI-FTICR-MS. DI-FTICR-MS provides molecular formulas in the absence of library matches for individual unknowns with significantly different abundances under drought conditions. Conversely, library matches from significantly changing classes of compounds may exemplify potential constituents. Molecular structure information gathered from library matches and confident molecular formula assignments can be extrapolated to unknown SOM components of significance when arranged into molecular networks with similar fragmentation spectra.

We also used library matches to validate molecular formula assignments. The number of library matches for SOM components was nearly doubled by searching GNPS libraries, which take advantage of open-source, community-populated tandem mass spectra to identify components. Matches with significantly different abundances between moisture conditions, though few, also increased. A comparable number of matches were found using our in-house library and the external GNPS libraries, yet, as expected, the coverage was different. Identifications from the in-house library were primarily *m/z* < 200 and included amino acids, nucleic acids and sugars while matches from external GNPS libraries included a broader range of masses and a greater diversity of natural products including secondary metabolites, polyphenols, terpenoids, carboxylic acids as well as common anthropogenic contaminants, which better reflected the diversity of SOM components. GNPS library searches also allowed for validation of more DI-FTICR-MS formula assignments (17 vs 9) with *m/z* > 200 than the in-house library alone, which was developed primarily for plant and microbial metabolites (Table S5c).

### 3.6 Future Perspectives

SOM chemistry is highly complex, and there are significant technical challenges in identifying its component parts. We demonstrate here that, GNPS networking, combined with ultra-high resolution mass accuracy formula assignments can assist with characterization and identification of SOM components. GNPS has also been used to discover natural products, primarily from plant and microbial organisms or consortia, and has the potential to aid in the discovery of novel SOM components [73]. As FTICR-MS, LC-MS/MS and molecular networking tools become more utilized for SOM, components found to be significant to carbon persistence in soils can be targeted for chromatographic separation, fraction collection, isolation, and identification. Ultimately, this will also improve libraries and can help reveal the pathways for SOM formation and endurance in soils.

The detritusphere represents a habitat with considerable microbial activity, where an abundance of carbon, including primary metabolites and other known compounds, are generated that have a greater likelihood of being matched to libraries than in bulk soils with less nutrients, microbial abundance, and arguably a greater proportion of carbon that has been modified from known compounds. As libraries are improved to include organic matter that has been modified from known precursors, such as lignin or cellulosic degradation products, this approach will become increasingly more useful for annotating SOM.

Although multiple orthogonal chromatographic separations may be required for a comprehensive evaluation of the structures and diversity of isomers, LC-MS-MS molecular networking integrated with direct infusion measurements, validates formula assignments, improves assignment rates and provides additional structural information for unknowns without sacrificing throughput and sensitivity (Figure 1). Our findings show that SOM composition changed significantly with drought in soil containing decaying organic matter. Average NOSC, O/C ratios and proportion of condensed aromatic compounds decreased while average H/C ratios and the proportion of lignin-like compounds increased. Putative matches for SOM components support these trends as well as the presence of fungal and bacterial metabolites. Reduced microbial activity along with reduced carbon decomposition and a shift in metabolite production due to changing microbial community composition or organisms under drought-stress are some of the underlying processes potentially driving these trends. However, most SOM components remain unidentified. Along with improved separations, expanded library databases for SOM and the potential for discovering novel compounds through networking, this integrated approach is a promising tool for improving annotation of SOM and understanding the molecular structures of carbon that persist in soils.

## Supporting Information

Supplemental methods, supporting information and supplemental tables and figures including molecular formula assignments, calculated average molecular properties, library matches, van Krevelen plots, LC-MS precursor mass chromatogram, selected example tandem mass spectra, and statistical analysis for lipid identifications.

## Acknowledgments

Funding for this work was provided by the USDOE Office of Science, Biological and Environmental Research, Biological Systems Science Division Grant # SCW1632. We would like to thank Montana Smith help with sample prep, Jason Toyoda for help with data collection and Peter Kennedy and the rest of the LLNL Soil Microbiome SFA for helpful discussion regarding fungal and microbial response to drought.

