## Supporting Information for "Improved characterization of soil organic matter by integrating FTICR-MS, liquid chromatography tandem mass spectrometry and molecular networking: a case study of root litter decay under drought conditions"

**Supplementary Methods:**

**Soil Samples:** Soil was collected from 0-10 cm depth (‘A’ mineral horizon) below a stand of *Avena barbata* at the University of California Hopland Research and Extension Center (39°00.106`N, 123°04.184`W). The site is a Mediterranean annual grassland ecosystem (MAT max/min = 23/7 C; MAP = 956 mm yr-1), where Avena spp. is the dominant vegetation. The soil is classified as a Typic Haploxeralfs of the Witherall-Squawrock complex, soil pH is ~5.6, contains 45% sand, 36% silt, and 19% clay. Initial C and N content were 2.2% and 0.24%, respectively.

Approximately 65 g of Hopland soil was mixed with *A. barbata* root litter fragments (1-5 mm) at a density of 0.013 g detritus dry g soil$^{-1}$ and packed into acrylic soil microcosms (56 × 56 × 76 linear cm) at field bulk density (1.21 g cm$^{3}$). Microcosms were placed inside growth chambers in a greenhouse with a 16-h light period per day (from 6 a.m. to 10 p.m.) and a maximum daytime and nighttime air temperature of 27°C and 24°C. Microcosms were maintained at one of two moisture treatments, to simulate differences in soil moisture during the spring growing season in California semiarid grasslands: ‘normal moisture’ (~16% ± 0.3 gravimetric soil moisture; mean ± standard error) or ‘drought’ conditions (~8% ± 0.5 gravimetric soil moisture). At 4, 8, and 12 weeks, microcosms were harvested from each treatment (n = 4) to collect soil samples.

**SOM Extraction**:
Step 1) A modified MPLEx procedure [1, 2] was performed on 1g each of lyophilized soil samples that were defrosted and added to a clean tube immediately prior to extraction. Water extractable carbon was removed first by adding 2 mL of milli-Q (<18.2 MΩ·cm resistivity) $H_2O$, shaking at 1000 rpm using a vortex shaker at room temperature for 2 hours, centrifuging at 6000 rpm for 5 minutes and removing the supernatant. This was repeated for the remaining soil pellet

[#]Current affiliation : Instituto Nacional de Investigación y Tecnología Agraria y Alimentaria: Madrid, Spain with another 2 mL of $H_2O$ and followed by 4 mL of chloroform, 2 mL of methanol and 0.25 mL of $H_2O$ for extraction of less polar compounds. The 2:1 chloroform: methanol slurry was shaken for 1 hour at 1000 rpm and incubated over night at 4 degrees C with another 1.25 mL of $H_2O$ to allow for bi-layer separation. Samples were centrifuged at 6000 rpm for 5 minutes prior to removal of each layer individually.

Step 2) Solid phase extraction (SPE) was performed for water extracts using Bond Elut PPL cartridges to remove salts and impurities that could interfere with MS analysis [3]. Water extracts were diluted to 5 mL with milliQ water and adjusted to pH 2 with 2uL of 85% $H_3PO_4$ prior to addition to PPL cartridges that had been activated with methanol. Organic matter (OM) bound to the cartridges was rinsed with 50 mL each of 10mM HCl, dried with nitrogen and eluted in 1.5 mL of methanol. Extracts were analyzed individually with DI-FTICR-MS and combined for LC-MS/MS analysis as follows:

Step 3) Drying: For each sample, 40 uL of methanol + 40 uL of chloroform from step 1 were dried in a speed vac.

Step 4) Reconstitution: Then, 40 uL of methanol that contained water-extracted SOM from step 2 was added to the vial from Step 3. No precipitation of OM due to reconstitution was observed for the samples in this study, likely due to the solvent exchange from water to methanol for water extracted OM and the combined "batch" nature of the extraction that included all three solvents simultaneously.

**DI-FTICR-MS:** Spectra over the *m/z* range 200-900 were optimized for peak intensity, shape, and resolution by adjusting the ion accumulation time for each sample within the range of 50-80 ms, 100-300 ms, and 50-100 ms for water, methanol and chloroform extracts, respectively. 1.1s transients were coadded over 144 acquisitions for an estimated average resolution of 240,000 at *m/z* 400. External calibration was performed using Agilent tune mix followed by internal recalibration of each spectrum, post-acquisition, with calibration lists of standard OM components. Peak picking was performed in Data Analysis using a signal to noise ratio (S/N) threshold of 7 and an absolute intensity threshold of 100,000 before exporting peak lists to Formularity for alignment using a 0.5 ppm threshold and formula assignment within a <0.5 ppm error (average error < 0.2 ppm)[4]. Elemental ratios were calculated for each molecular formula and averaged for each sample. The nominal oxidation state of carbon was calculated for each molecular formula according to the following [5]:

(1) $$NOSC = -\left(\frac{-Z+4a+b-3c-2d+5e-2f}{a}\right)+4$$

Where Z is the net charge and a, b, c, d, e and f are the number of atoms of C, H, N, O, P and S, respectively for each molecular formula. Here, Z= -1, S=0 and P=0 and the equation reduces to:

(2) $$NOSC = -\left(\frac{4C+H-3N-2O}{C}\right)+4$$

**Metabolite Chromatography:** At a flow rate of 0.3 mL/min, the elution gradient started stable at 90% A (10% B) for 5 minutes before it linearly changed to 10% A (90%B) during the 15 consecutive minutes (min 20) and then maintained constant for 2 minutes (min 22). The initial conditions were linearly recovered over an additional 2 minutes (min 24). Washing and stabilization of the column was performed at the initial conditions for 11 additional minutes prior to the next injection (min. 35).

**Lipid Chromatography:** Chloroform-only extracts were analyzed for lipids as outlined by Kyle et al. [6] using a Waters HPLC system containing a reversed phase C18 CSH column (150×3 mm, 1.7 µm particle size) coupled to a LTQ Orbitrap Velos high-resolution mass spectrometer for analysis. After injection of 10 uL per sample, lipid molecular species were separated over a 34-minute gradient. LC mobile phases consisted of ACN/H2O (40:60) containing 10 mM ammonium acetate (A) and ACN/IPA (10:90) containing 10 mM ammonium acetate (B) conducted at a flow rate of 250 µl/min. The elution gradient is provided in Table S3 of Kyle et al. [6] Washing and stabilization of the column was performed at the initial conditions for 8 minutes prior to the next injection.

**High Resolution Mass Spectrometry (HRMS) Measurements for Metabolites:** HESI was performed with a capillary voltage of 3.5kV in negative mode and 4kV in positive mode with a sheath gas flow of 50, auxiliary gas flow of 30 and sweep gas flow of 2 at 300C. For each ionization mode (positive and negative), the HRMS operated at full-scan mode, high resolving power (60,000 at *m/z* 400) with an AGC target of 30,000, and tandem mass spectrometry AGC target of 10,000 to detect ions between *m/z* 70 and 800. Data dependent acquisition (DDA) was utilized to select the top 10 most intense precursor ions in each scan for fragmentation in the ion trap, without dynamic exclusion enabled. Acquisition speed averaged 2.4 s per spectrum.

**HRMS Measurements for Lipids** A heated electrospray ionization source (HESI) with a capillary voltage of 3.5 kV in negative mode and 3.5 kV in positive mode and gas flows of 45, 30, and 2 (sheath, auxiliary, sweep, respectively) was used to generate ions for HRMS analysis. HRMS data were collected separately in both positive and negative full-scan mode and high resolving power (120,000 at *m/z* 400) with an AGC target of 1000000 and tandem mass spectrometry AGC target of 10000 for ion trap MSn and 50000 for FTMS MSn to detect ions between *m/z* 200-2000. Acquisition speed averaged 0.24 s per scan, and DDA was used to select the top 8 most intense precursor ions peaks in each spectrum for fragmentation in the ion trap. Lipids were fragmented using both higher-energy collision dissociation (HCD) and collision-induced dissociation (CID). QC checks were performed with Avanti Lipids Porcine Brain Total Lipid Extract standards at a 0.5mg/mL concentration, with two blanks for every 30 samples. Spectra for each ionization mode were aligned and gap filled separately using MZmine and LIQUID for confident lipid identifications[6]. Aligned features were manually verified and peak apex intensity values for identified lipids were normalized to the total intensity of the chromatogram for each sample prior to export. Statistical analysis was performed using MetaboAnalyst 5.0, for which details are provided below [7].

**Metabolite Data Processing:** Baseline chromatograms were corrected for each sample individually and metabolite features with unique masses and retention times (RTs) were detected, deconvoluted, aligned and gap filled according to parameters listed in Table S2, before matching against an in-house constructed metabolite library, based on RT with an allowed shift of ≤ 0.2 min and the monoisotopic exact mass of standards with an error of ≤ 7ppm. Peak areas were subsequently exported to a CSV file. Peak areas were normalized to the total intensity of the chromatography of each sample prior to statistical analysis to account for run-to-run variation and assess differences between soil metabolites between drought and control over the three time points (4, 8 and 12 weeks).

**Statistical analysis of lipids using MetaboAnalyst:** Identified lipids and their relative abundances from negative mode spectra only for each sample treatment replicate were uploaded, log transformed and scaled by mean centering using the MetaboAnalysis one-factor statistical analysis module. Pairwise score plots were generated for 5 PCs, of which PC1 and PC2 explained 67.2 and 16.5% of the variance, respectively. A 2D scores plot was generated for PC1 and PC2 displaying the 95% confidence regions (Figure S7a). Normalized data was also used to create a heatmap showing the top 25 lipids with the most significant difference in abundances in the drought compared to the control, based on a T-test/ANOVA, Euclidian distance measure and Ward cluster algorithm.

**Quality of GNPS matches:** A few matches to GNPS library spectra were catalogued as positive mode adducts and therefore removed from this analysis. Tryptophan and gluconic acid were two exceptions in which the negative mode m/z was appropriate and supported by in-house library matches. Occasionally the same *m/z* was matched to conflicting compounds from each library (Table S5c-d). *m/z* 269.046 with RT 16.98 minutes matched to the GNPS library spectrum for aloe-emodin with just three fragment ion peaks and the same *m/z* with RT 17.41 min was matched to our in-house library standard emodin, a structural isomer of aloe-emodin differing by a hydroxyl and methyl group instead of a hydroxy methyl group. Though the difference in retention times suggest the possibility both isomers could be present, this is not conclusive. A diagnostic peak (240 *m/z*), used to discriminate negative mode spectra of aloe-emodin from emodin [8] was detected in the experimental spectra as well as both GNPS library spectra for emodin and aloe-emodin, thus allowing for conflicting matches (Figure S5a). However, this fragment ion was not considered in the GNPS library matching algorithm, instead three other ions (*m/z* 225, 197 and 181) were used to match the experimental spectra to the GNPS library spectrum, all of which were also present in the GNPS library spectra for both emodin and aloe-emodin. GNPS generates a consensus spectrum from a group of individual raw tandem mass spectra scans that are not always representative of the individual spectra, and slight differences in averaged *m/z* or intensities can increase or decrease the cosine similarity score to a library spectrum, thus excluding a potential match depending on the search threshold. Fragmentation spectral quality can also vary with the type of instrument, collision energy, and mechanisms of fragmentation, making comparisons incomplete and resulting in incorrect matching in some cases or lack of matches in others. Usnic acid is another GNPS library match that was also verified by our in-house library standard. 4 fragment ion peaks were used to match to the GNPS library, yet upon further inspection of individual scans, up to 7 fragment ions could be used to verify this library match

**Biological Relevance of Library Matches- Support from Lipidomics:** 100 lipids were identified using our in-house library from negative ion mode spectra and 202 lipids were identified from positive mode spectra that provide evidence for potential community members present in the detritus soils. Over 1/3 of the total lipids (121) detected across both treatments were triacylglycerolipids (TG), which are known to be highly abundant in fungi [9]. Overall, statistical analysis of in-house lipid identifications supported DI-FTICR-MS and metabolomics results from harvest 3 (week 12) showing lipids under drought in the detritusphere soil were statistically different from the control (Figure S7a). A heatmap of the top 25 lipids from both positive and negative mode spectra with the most significantly different abundances between the drought and control treatments ($P<0.08$) after 12 weeks is shown in Figure S7b. Lipids with the most significant difference in abundance ($P<0.05$, Figure 7b) include lipids from bacteria that make phosphatidylethanolamine (PE) odd-chain lipids and contain odd chains with one ring or double bond (based on the *m/z* or the fragment ion). The latter indicates a cyclopropanyl ring may be present, which is known to be produced in response to environmental stress [10]. Other lipids (i.e. PE(P_15:0:15:0)) with significantly higher abundance ($P<0.05$) in the drought may come from anaerobes, as indicated by the presence of plasmalogens, an ether lipid sensitive to the presence of oxygen and signifying cell resistance to adverse environmental conditions [11]. Eucaryotic phosphatidylcholine (PC) lipids containing 20:4 chains (likely the fatty acid arachidonic acid) were found to be less abundant under drought conditions, while lipids produced by eucaryotic photosynthetic organisms or plant species were found to be significantly more abundant in the drought, including PE lipids containing 20:3 chains (possibly dihomo-gamma-linolenic acid), PE and phosphatidylglycerol (PG) containing 18:3 chains (likely the fatty acid linolenic acid), and triacylglycerol (TG) lipids ($P<0.05$) [12-14]. Additionally, DGDG (16:0/19:1) chloroplast lipids that contain 1 double bond or cyclopropanyl ring suggesting the organism is under stress were also detected at significantly higher abundance under drought conditions. Lipids indicative of eucaryotic photosynthesis could include lichen, or close fungal relatives to the lichen fungal partner. The presence of these organisms is supported by the detection of metabolites specific to lichen that were matched to GNPS libraries.

Greater than 80% of the in-house lipidomics matches were for *m/z* > 500, thus, there was less commonality between in-house lipid identifications (precursor masses with tandem mass spectra), and *m/z* detected with direct infusion FTICR-MS for the chloroform extract alone. However, chloroform extracts were also analyzed for matches to secondary metabolites and other compounds contained in GNPS libraries. Several additional precursor masses were matched to GNPS libraries using lipidomics datasets, including lichen- and fungi- specific secondary metabolites neodihydroprotolichesterinic acid (*m/z* 325.240), chloratranorin*m/z* (*m/z* 407.053) and soyacerebrocide I (*m/z* 712.539). The precursor mass, chloratranorin, shared an edge with atranorin (*m/z* 373.094, (also detected and matched to the GNPS library using the metabolomics datasets) which differs in structure from it only by a hydrogen in place of the chlorine.

### Supplementary Tables and Figures

Table S1. Measured carbon values for drought and control samples
Table S2. MZmine LC-MS metabolomics processing parameters
Table S3. GNPS Classical molecular networking parameters
Table S4. Molecular formula assignments and calculated properties
Table S5a-d. In-house and GNPS library matches
Table S6. Precursor masses from polyphenol subnetwork (Figure 5b) matched to GNPS or detected with DI-FTICR-MS

Figure S1. van Krevelen plots of drought vs control detritusphere SOM at week 12
Figure S2. van Krevelen plot of FTICR-MS molecular formulas also detected with LC-MS at significantly higher abundance in the drought
Figure S3. LC vs RT chromatogram of precursor masses overlayed with DI-FTICMRS masses and formulas
Figure S4. Venn diagram of DI-FTICRMS *m/z* detected as precursor *m/z* with LC-MS/MS spectra and assigned molecular formula or matched to GNPS libraries.
Figures S5a-i. Tandem mass spectra and mirror match spectra for aloe-emodin, emodin, usnic acid, and GNPS aligned spectra for endocrocin, *m/z* 363.036 and cis-5,8,11-Eicosatrienoic acid with aloe-emodin.
Figure S6. van Krevelen diagrams for drought only and control only CHO and CHON molecular formalas
Figure S7. PCA and heatmap for lipids identified from detritusphere soil extracts at 12 weeks

GNPS Classical Molecular Networking Jobs

Metabolomics Negative mode, Detritusphere Harvest 3:
https://gnps.ucsd.edu/ProteoSAFe/status.jsp?task=499f3454320146b0ac6caba3b0486ef4

Metabolomics Negative mode, Detritusphere Harvest 1-3:
https://gnps.ucsd.edu/ProteoSAFe/index.jsp?task=b99361332cc0460f9e8190407d405f04

Metabolomics Negative mode, All Treatments, Harvest 1-3:
https://gnps.ucsd.edu/ProteoSAFe/status.jsp?task=7f4122b7fa9145fda6ef9c2b0f21be2e

Table S1. Average TOC (%C) Values for Drought and Control Detritusphere Soils

| Weeks | 4 | | 8 | | 12 | |
|---|---|---|---|---|---|---|
| Moisture | Control | Drought | Control | Drought | Control | Drought |
| TOC mean | 2.25 | 2.38 | 2.58 | 2.29 | 2.23 | 2.26 |
| TOC stdev | 0.10 | 0.15 | 0.65 | 0.08 | 0.16 | 0.09 |

Table S2. Metabolomics MZmine Processing Parameters

| | | |
|---|---|---|
| 1 | Baseline correction – RollingBall baseline corrector | |
| | Chromatogram type | TIC |
| | Use m/z bins | No |
| | wm | 20 |
| | ws | 20 |
| 2 | Mass detection (exact Mass) | |
| | Noise level | 1000 |
| 3 | FTMS shoulder peaks filter | |
| | Peak model function | Loretzian |
| | Mass Resolution | 60,000 |
| 4 | ADAP Chromatogram builder | |
| | Minimum size in number of scans | 5 |
| | Group intensity threshold | 1000 |
| | Minimum highest intensity | 5000 |
| | m/z tolerance | 0.0005 or 7ppm |
| 5 | Smoothing | |
| | Filter width | 9 |
| 6 | Chromatogram deconvolution (local minimum search) | |
| | Chromatographic threshold | 95% |
| | Search minimum in RT range (min) | 0.3 |
| | Minimum relative height | 30% |
| | Minimum absolute height | 5000 |
| | Minimum ratio of peak top/edge | 1.3 |
| | Peak duration range | 0-2 min |
| 7 | Chromatogram alignment (join alignment) | |
| | m/z tolerance | 0.0005 or 7ppm |
| | Weight for m/z | 60 |
| | RT tolerance | 0.25 min |
| | Weight for RT | 40 |
| 8 | Gap filling (Peak Finder) | |
| | Intensity tolerance | 3% |
| | m/z tolerance | 0.0005 or 6ppm |
| | Retention time tolerance | 0.2 min |
| | RT correction | No |
| 9 | Filtering (Duplicate Peak Filter) | |
| | Filter mode | New Average |
| | m/z tolerance | 0.0005 or 7ppm |
| | RT tolerance | 0.2 min |
| 10 | Metabolite Assignation (Custom Database) | |
| | m/z tolerance | 0.001 or 15ppm* |
| | RT tolerance | 0.5 min* |
| 11 | Data Exported | Peak Area |

Table S3. GNPS Classical Molecular Networking Parameters

| Basic Options | Value |
|---|---|
| Precursor Mass Tolerance | 0.02 Da |
| Fragment Ion Mass Tolerance | 0.5 Da |
| Network Options | |
| Network Min Cosine Score for Pairs | 0.7 |
| Network TopK | 10 |
| Maximum Connected Component Size | 100 |
| Minimum Matched Fragment Ions | 3 |
| Minimum Cluster Size | 2 |
| Library Search Options | |
| Minimum Matched Peaks | 3 |
| Search Analogs | No |
| Score Threshold | 0.7 |

Table S4. FTICR-MS Average Assignments and Calculated Properties for Water, Methanol and Chloroform Extractable Organic Matter

| Weeks | 4 | | | 8 | | | 12 | |
|---|---|---|---|---|---|---|---|---|
| Moisture | Control | Drought | | Control | Drought | | Control | Drought |
| Mean H2O Peak No. | 4961 | 5765 | | 5752 | 5218 | | 5791 | 5373 |
| Mean Assignments | 2869 | 3513 | | 3485 | 3122 | | 3761 | 3239 |
| Mean % Assigned N<=2 SP =0 | 58% | 61% | | 61% | 60% | | 65% | 60% |
| Monoisotopic_Assignments | 2406 | 2842 | | 2836 | 2510 | | 3037 | 2637 |
| Isotopologue_Assignments | 464 | 671 | | 649 | 612 | | 724 | 603 |
| Mean_Error_PPM | 0.114 | 0.120 | | 0.118 | 0.117 | | 0.123 | 0.119 |
| DBE_Mean | 9.81 | 10.04 | | 10.09 | 9.79 | | 10.28 | 9.88 |
| AI_Mean | 0.103 | 0.110 | | 0.113 | 0.106 | | 0.114 | 0.123 |
| AImod_Mean | 0.309 | 0.314 | | 0.322 | 0.303 | | 0.324 | 0.308 |
| NOSC_Mean | -0.04 | -0.02 | | 0.00 | -0.06 | | 0.01 | -0.07 |
| NOSC_median | -0.01 | 0.00 | | 0.01 | -0.04 | | 0.01 | -0.07 |
| OC_Mean | 0.530 | 0.529 | | 0.531 | 0.522 | | 0.534 | 0.516 |
| HC_Mean | 1.147 | 1.134 | | 1.120 | 1.154 | | 1.115 | 1.150 |

| Weeks | 4 | | | 8 | | | 12 | |
|---|---|---|---|---|---|---|---|---|
| Moisture | Control | Drought | | Control | Drought | | Control | Drought |
| Mean MeOH Peak No. | 2990 | 2567 | | 3580 | 3199 | | 2440 | 3467 |
| Mean Assignments | 1758 | 1696 | | 1962 | 2101 | | 1612 | 2060 |
| Mean % Assigned N<=2 SP =0 | 59% | 66% | | 55% | 66% | | 66% | 59% |
| Monoisotopic_Assignments | 1522 | 1491 | | 1692 | 1786 | | 1437 | 1769 |
| Isotopologue_Assignments | 236 | 205 | | 270 | 315 | | 175 | 292 |
| Mean_Error_PPM | 0.087 | 0.083 | | 0.092 | 0.094 | | 0.085 | 0.097 |
| DBE_Mean | 6.73 | 6.82 | | 7.08 | 7.37 | | 6.89 | 7.48 |
| AI_Mean | 0.125 | 0.127 | | 0.150 | 0.142 | | 0.147 | 0.136 |
| AImod_Mean | 0.188 | 0.196 | | 0.207 | 0.216 | | 0.197 | 0.218 |
| NOSC_Mean | -0.58 | -0.58 | | -0.54 | -0.53 | | -0.55 | -0.51 |
| NOSC_median | -0.59 | -0.58 | | -0.53 | -0.52 | | -0.55 | -0.51 |
| OC_Mean | 0.437 | 0.432 | | 0.438 | 0.434 | | 0.441 | 0.440 |
| HC_Mean | 1.461 | 1.446 | | 1.426 | 1.406 | | 1.441 | 1.399 |

| Weeks | 4 | | | 8 | | | 12 | |
|---|---|---|---|---|---|---|---|---|
| Moisture | Control | Drought | | Control | Drought | | Control | Drought |
| Mean CHCl3 Peak No. | 2335 | 2541 | | 2312 | 1939 | | 2172 | 2058 |
| Mean Assignments | 1183 | 1392 | | 1230 | 1086 | | 1085 | 803 |
| Mean % Assigned N<=2 SP =0 | 51% | 55% | | 53% | 56% | | 50% | 39% |
| Monoisotopic_Assignments | 826 | 950 | | 859 | 760 | | 766 | 597 |
| Isotopologue_Assignments | 358 | 442 | | 371 | 326 | | 319 | 206 |
| Mean_Error_PPM | 0.16 | 0.17 | | 0.16 | 0.16 | | 0.16 | 0.16 |
| DBE_Mean | 4.24 | 4.47 | | 4.53 | 4.13 | | 4.67 | 5.21 |
| AI_Mean | 0.06 | 0.05 | | 0.06 | 0.05 | | 0.08 | 0.10 |
| AImod_Mean | 0.08 | 0.08 | | 0.08 | 0.07 | | 0.09 | 0.09 |
| NOSC_Mean | -1.40 | -1.38 | | -1.37 | -1.41 | | -1.36 | -1.32 |
| NOSC_median | -1.39 | -1.38 | | -1.37 | -1.41 | | -1.36 | -1.32 |
| OC_Mean | 0.186 | 0.190 | | 0.191 | 0.186 | | 0.194 | 0.203 |
| HC_Mean | 1.77 | 1.77 | | 1.75 | 1.79 | | 1.75 | 1.73 |

Table S5a. In-house Library Matches (Negative and Positive Mode)

| In-house Library Match | m/z | RT (min) | Mode | Library | P_value | Abundance Significance (P<0.05) |
|---|---|---|---|---|---|---|
| Glycine | 76.0393 | 1.31 | POS | In-house | 0.28341 | |
| Pyruvic.acid | 87.0088 | 1.51 | NEG | In-house | 0.12495 | |
| GLYCERALDEHYDE.Peak.1.2 | 89.0244 | 1.39.1.77 | NEG | In-house | 0.55338 | |
| Alanine | 90.0550 | 1.3 | POS | In-house | 0.00000 | Drought |
| Aniline | 94.0651 | 1.93 | POS | In-house | 0.11109 | |
| SUCCINATE.SEMIALDEHYDE | 101.0244 | 1.79 | NEG | In-house | 0.43568 | |
| AMINOCYCLOPROPANE.1.CARBOXYLATE | 102.0550 | 1.29 | POS | In-house | 0.04810 | Drought |
| Hydroxybutyric.acid.Peak.2 | 103.0401 | 1.945 | NEG | In-house | 0.08987 | |
| L.SERINE | 104.0353 | 1.29 | NEG | In-house | 0.42644 | |
| AminoIsoButyric.acid | 104.0706 | 1.31 | POS | In-house | 0.19120 | |
| DIETHANOLAMINE | 106.0863 | 1.24 | POS | In-house | 0.24032 | |
| CYTOSINE | 112.0505 | 1.26 | POS | In-house | 0.05735 | |
| URACIL | 113.0346 | 1.37 | POS | In-house | 0.69901 | |
| DIHYDROURACIL | 113.0357 | 1.355 | NEG | In-house | 0.27991 | |
| MALEAMATE | 116.0342 | 1.44 | POS | In-house | 0.47125 | |
| L.PROLINE | 116.0706 | 1.35 | POS | In-house | 0.60821 | |
| N.ACETYLGLYCINE. | 118.0499 | 1.55 | POS | In-house | 0.06923 | |
| L.THREONINE | 118.0510 | 1.31 | NEG | In-house | 0.40023 | |
| L.VALINE | 118.0863 | 1.35 | POS | In-house | 0.07497 | |
| 5.AMINOPENTANOATE | 118.0863 | 1.28 | POS | In-house | 0.06369 | |
| L.VALINE | 118.0863 | 1.35 | POS | In-house | 0.12596 | |
| PURINE.Peak.1 | 119.0363 | 1.38 | NEG | In-house | 0.03397 | Drought |
| PICOLINIC.ACID | 124.0393 | 1.35 | POS | In-house | 0.95507 | |
| Phloroglucinol | 127.0390 | 1.43 | POS | In-house | 0.55746 | |
| THYMINE | 127.0502 | 1.39.1.85 | POS | In-house | 0.11182 | |
| OXO.L.PROLINE.Peak.1 | 128.0353 | 1.37 | NEG | In-house | 0.25125 | |
| METHYL.2.OXO.PENTANOIC.ACID | 129.0557 | 5.74 | NEG | In-house | 0.25585 | |
| L.PIPECOLIC.ACID | 130.0863 | 1.35.1.48 | POS | In-house | 0.41613 | |
| LEUCINE.Peak.1.2 | 130.0874 | 1.845 | NEG | In-house | 0.04769 | Drought |
| L.ASPARAGINE | 131.0462 | 1.31 | NEG | In-house | 0.18108 | |
| L.ORNITHINE | 131.0826 | 1.245 | NEG | In-house | 0.20336 | |
| ASPARTATE | 132.0302 | 1.41 | NEG | In-house | 0.60625 | |
| ADENINE | 134.0472 | 1.345 | NEG | In-house | 0.04885 | Normal |
| SALICYLATE | 137.0244 | 11.39 | NEG | In-house | 0.74407 | |
| UROCANATE | 137.0357 | 1.325 | NEG | In-house | 0.54316 | |
| Hypoxanthine | 137.0458 | 1.4 | POS | In-house | 0.00266 | Normal |
| p.Anisaldehyde | 137.0597 | 12.72 | POS | In-house | 0.82681 | |
| TRIGONELLINE | 138.0550 | 1.36 | POS | In-house | 0.26838 | |

| | | | | | | |
|---|---|---|---|---|---|---|
| L.GLUTAMINE | 145.0619 | 1.315 | NEG | In-house | 0.92714 | |
| L.GLUTAMIC.ACID | 146.0459 | 1.32 | NEG | In-house | 0.01547 | Drought |
| GUANINE | 152.0567 | 1.33 | POS | In-house | 0.02080 | Normal |
| L.HISTIDINE | 154.0622 | 1.255 | NEG | In-house | 0.65496 | |
| 3.HYDROXY.3.METHYLGLUTARATE | 161.0456 | 1.845 | NEG | In-house | 0.02128 | Drought |
| N.METHYL.L.GLUTAMATE | 162.0761 | 1.34 | POS | In-house | 0.14913 | |
| L.CARNITINE | 162.1125 | 1.27 | POS | In-house | 0.30139 | |
| HYDROXY.3.METHYLGLUTARATE | 163.0601 | 1.85 | POS | In-house | 0.01105 | Drought |
| Eugenol | 163.0764 | 14.61 | NEG | In-house | 0.37297 | |
| L.PHENYLALANINE.Peak.1.2 | 164.0717 | 2.185 | NEG | In-house | 0.74209 | |
| L.PHENYLALANINE | 166.0863 | 2.19 | POS | In-house | 0.35929 | |
| 3.DEHYDROSHIKIMATE | 171.0299 | 1.45 | NEG | In-house | 0.08145 | |
| SUBERIC.ACID | 173.0819 | 9.84 | NEG | In-house | 0.64230 | |
| Citrulline | 174.0884 | 1.285 | NEG | In-house | 0.07878 | |
| L.ARGININE | 175.1190 | 1.25 | POS | In-house | 0.98066 | |
| D.GULONIC.ACID.GAMA.LACTONE | 177.0405 | 1.355 | NEG | In-house | 0.66163 | |
| Methyleugenol | 177.0921 | 16.095 | NEG | In-house | 0.27340 | |
| Sugars.Monosaccharides.Hexoses | 179.0561 | 1.31 | NEG | In-house | 0.12823 | |
| Sugars.Alcohol.Hexoses | 181.0717 | 1.31 | NEG | In-house | 0.02509 | Drought |
| L.TYROSINE | 182.0812 | 1.37 | POS | In-house | 0.09603 | |
| EPINEPHRINE | 184.0968 | 1.35 | POS | In-house | 0.09205 | |
| AZELAIC.ACID | 187.0976 | 11.3 | NEG | In-house | 0.34514 | |
| HYDROXYDECANOATE | 187.1340 | 12.7 | NEG | In-house | 0.83562 | |
| NALPHA.ACETYL.L.LYSINE | 189.1234 | 1.28 | POS | In-house | 0.02592 | Drought |
| GLUCONIC.ACID | 195.0510 | 1.52 | NEG | In-house | 0.56646 | |
| GLUCOSAMINATE | 196.0816 | 1.32 | POS | In-house | 0.00752 | Drought |
| Tryptophan | 205.0972 | 3.5 | POS | In-house | 0.30054 | |
| Jasmonic.acid | 211.1329 | 13.47 | POS | In-house | 0.23540 | |
| O.SUCCINYL.L.HOMOSERINE | 220.0816 | 1.36 | POS | In-house | 0.19253 | |
| THYMIDINE.Peak.2 | 241.0830 | 1.96 | NEG | In-house | 0.93616 | |
| Cytidine | 242.0783 | 1.29 | NEG | In-house | 0.09012 | |
| DEOXYADENOSINE | 252.1091 | 1.36.1.82 | POS | In-house | 0.84777 | |
| PALMITOLEIC.ACID | 253.2173 | 23.015 | NEG | In-house | 0.62557 | |
| ADENOSINE.Peak.1.2 | 266.0895 | 1.38 | NEG | In-house | 0.01076 | Drought |
| Inosine.Peak.1 | 267.0735 | 1.41 | NEG | In-house | 0.05600 | |
| Emodin | 269.0455 | 17.41 | NEG | In-house | 0.00001 | Drought |
| GAMMA.LINOLENIC.ACID | 279.2319 | 19.34.18.1 | POS | In-house | 0.32937 | |
| LINOLEATE | 281.2475 | 17.48.22.48 | POS | In-house | 0.02984 | Drought |
| RAC.GLYCEROL.1.MYRISTATE | 303.2530 | 21.31 | POS | In-house | 0.03462 | Normal |
| Sugars.Disaccharides | 341.1089 | 1.345 | NEG | In-house | 0.22951 | |
| Usnic.acid | 343.0823 | 20.455 | NEG | In-house | 0.08421 | |

| | | | | | | |
|---|---|---|---|---|---|---|
| Rhapontin | 421.1493 | 10.64 | POS | In-house | 0.01553 | Drought |
| Sugars.Trisaccharides | 503.1617 | 1.29 | NEG | In-house | 0.08278 | |

Table S5b. GNPS Library Matches to Negative Mode Spectra

| GNPS Library Match | GNPS m/z | Adduct | RT mean (min) | Shared Peaks | Cosine Score | Library | P_value | Abundance Significance (P<0.05 ) |
|---|---|---|---|---|---|---|---|---|
| KO000888 Gallate\|Pyrogallol-5-carboxylic acid\|3,4,5-Trihydroxybenzoate\|3,4,5-Trihydroxybenzoic acid\|Gallic acid | 169.000 | [M-H]- | 1.80 | 4 | 0.75 | GNPS/ Massbank | 0.06603 | |
| Juarezic Acid | 173.046 | [M-H]- | 21.04 | 4 | 0.71 | GNPS | | |
| RP015311 Suberic acid\|Octanedioic acid | 173.082 | [M-H]- | 9.50 | 4 | 0.98 | GNPS/ Massbank | 0.64230 | |
| esculetin\|6,7-dihydroxychromen-2-one | 177.019 | [M-H]- | 20.12 | 3 | 0.92 | GNPS/ Massbank | 0.00988 | Drought |
| 3-Amino-Beta-Pinene | 186.114 | [M-H]- | 8.86 | 3 | 0.82 | GNPS | | |
| AZELAIC ACID | 187.098 | [M-H]- | 10.78 | 4 | 0.92 | GNPS | 0.34514 | |
| KNA00638 D-Gluconic acid\|D-Gluconate\|D-gluco-Hexonic acid | 195.051 | M+H | 1.43 | 6 | 0.91 | GNPS/ Massbank | 0.56646 | |
| KO001812 Syringate\|Syringic acid | 197.009 | [M-H]- | 4.36 | 3 | 0.80 | GNPS/ Massbank | | |
| SEBACATE | 201.113 | [M-H]- | 12.17 | 3 | 0.79 | GNPS | 0.36816 | |
| KNA00633 L-Tryptophan\|Tryptophan\|(S)-alpha-Amino-beta-(3-indolyl)-propionic acid | 203.083 | M+H | 3.38 | 7 | 1.00 | GNPS/ Massbank | | |
| LU047654 Undecanedioic acid | 215.129 | [M-H]- | 13.21 | 4 | 0.72 | GNPS/ Massbank | | |
| Spectral Match to Leu-Val from NIST14 | 229.156 | [M-H]- | 2.55 | 5 | 0.85 | GNPS | 0.05048 | |
| THYMIDINE | 241.083 | [M-H]- | 1.85 | 4 | 0.79 | GNPS | 0.93616 | |
| (Z)-2-octylpent-2-enedioic acid | 241.145 | [M-H]- | 13.93 | 3 | 0.78 | GNPS | | |
| (4aR,5S,8aS,9aR)-9a-hydroxy-3,4a,5-trimethyl-5,6,7,8,8a,9-hexahydro-4H-benzo[f][1]benzofuran-2-one | 249.15 | [M-H]- | 20.45 | 5 | 0.89 | GNPS | 0.00002 | Drought |
| Spectral Match to Dodecyl sulfate from NIST14 | 265.148 | [M-H]- | 24.32 | 7 | 0.92 | GNPS | | |
| DEOXYGUANOSINE | 266.089 | [M-H]- | 1.46 | 4 | 0.88 | GNPS | 0.01076 | Drought |
| LU108653 Aloe-emodin | 269.046 | [M-H]- | 16.98 | 3 | 0.79 | GNPS/ Massbank | 0.00001 | Drought |
| 3-(2,6-dihydroxyphenyl)-4-hydroxy-6-methyl-3H-2-benzofuran-1-one | 271.061 | [M-H]- | 12.80 | 5 | 0.71 | GNPS | 0.78298 | |
| Spectral Match to Mono-2-ethylhexyl phthalate from NIST14 | 277.145 | [M-H]- | 18.71 | 11 | 0.97 | GNPS | | |
| Spectral Match to Pinolenic acid from NIST14 | 277.217 | [M-H]- | 20.49 | 6 | 0.83 | GNPS | | |
| 3',4',5,7-tetrahydroxyflavone | 285.041 | [M-H]- | 16.08 | 6 | 0.80 | GNPS | | |
| 3',4',5,7-tetrahydroxyflavone | 285.065 | [M-H]- | 14.80 | 3 | 0.73 | GNPS | | |
| Spectral Match to Pregna-4,16-diene-3,20-dione from NIST14 | 293.176 | M-H2O-H | 18.77 | 4 | 0.73 | GNPS | 0.13246 | |
| Spectral Match to 5-HETE from NIST14 | 301.217 | N/A | 21.52 | 7 | 0.93 | GNPS | 0.15320 | |

| | | | | | | | | |
|---|---|---|---|---|---|---|---|---|
| Spectral Match to 5S-Hydroxy-6E,8Z,11Z-eicosatrienoic acid from NIST14 | 303.233 | M-H-H2O | 22.61 | 3 | 0.92 | GNPS | 0.95453 | |
| Spectral Match to cis-5,8,11-Eicosatrienoic acid from NIST14 | 305.248 | [M-H]- | 23.75 | 5 | 0.79 | GNPS | | |
| 1-[2-methyl-6-[(2S,3R,4S,5S,6R)-3,4,5-trihydroxy-6-(hydroxymethyl)oxan-2-yl]oxyphenyl]ethanone | 311.169 | [M-H]- | 19.22 | 3 | 0.72 | GNPS | 0.00613 | Drought |
| Endocrocin | 313.035 | [M-H] | 12.88 | 3 | 0.84 | GNPS | | |
| SM876251 Dodecylbenzenesulfonic acid | 325.185 | [M-H]- | 19.62 | 5 | 0.76 | GNPS/ Massbank | | |
| Spectral Match to D-(+)-Trehalose from NIST14 | 341.109 | [M-H]- | 1.32 | 7 | 0.97 | GNPS | 0.22951 | |
| BML00262 Usnic acid | 343.082 | [M-H]- | 19.71 | 4 | 0.82 | GNPS/ Massbank | 0.08421 | |
| Atranorin | 373.093 | [M-H]- | 20.11 | 7 | 0.74 | GNPS | 0.30345 | |
| NCGC00380423-01!2-(hydroxymethyl)-5-(2-oxopropyl)-7-[(2S,3R,4S,5S,6R)-3,4,5-trihydroxy-6-(hydroxymethyl)oxan-2-yl]oxychromen-4-one | 455.102 | M+FA-H | 1.33 | 4 | 0.75 | GNPS | | |
| (4aS,6aS,6bR,9R,10R,11R,12aR)-10,11-dihydroxy-9-(hydroxymethyl)-2,2,6a,6b,9,12a-hexamethyl-1,3,4,5,6,6a,7,8,8a,10,11,12,13,14b-tetradecahydropicene-4a-carboxylic acid | 487.343 | [M-H]- | 15.92 | 3 | 0.73 | GNPS | | |
| BML00943 Animicin A | 547.266 | M-H | 21.51 | 8 | 0.76 | GNPS/ Massbank | | |

*N/A = not annotated in GNPS library

Table S5c. Combined Library Matches from Negative Ion Mode Spectra (LC-MS, GNPS and DI-FTICR-MS)

| Library | In-house m/z | GNPS m/z | In-house RT (min) | GNPS RT mean (min) | GNPS Adduct | In-house Library Match | GNPS Library Match | DI-FTICR-MS m/z | DI-FTICR-MS Assigned Formula | DI-FTICR-MS Error (ppm) | van Krevelen Elemental Class | P_value | P<0.05 and treatment with higher abundance |
|---|---|---|---|---|---|---|---|---|---|---|---|---|---|
| In-house | 87.0088 | | 1.5 | | NEG | Pyruvic.acid | | ND | | | | 0.12495 | |
| In-house | 89.0244 | | 1.4 | | NEG | GLYCERALDEHYDE.Peak.1.2 | | ND | | | | 0.55338 | |
| In-house | 101.0244 | | 1.8 | | NEG | SUCCINATE.SEMIALDEHYDE | | ND | | | | 0.43568 | |
| In-house | 103.0401 | | 1.9 | | NEG | X2.Hydroxybutyric.acid.Peak.2 | | ND | | | | 0.08987 | |
| In-house | 104.0353 | | 1.3 | | NEG | L.SERINE | | ND | | | | 0.42644 | |
| In-house | 113.0357 | | 1.4 | | NEG | X5.6.DIHYDROURACIL | | ND | | | | 0.27991 | |
| In-house | 118.0510 | | 1.3 | | NEG | L.THREONINE | | ND | | | | 0.40023 | |
| In-house | 119.0363 | | 1.4 | | NEG | PURINE.Peak.1 | | ND | | | | 0.03397 | Drought |
| In-house | 128.0353 | | 1.4 | | NEG | X5.OXO.L.PROLINE.Peak.1 | | ND | | | | 0.25125 | |
| In-house | 129.0557 | | 5.7 | | NEG | X4.METHYL.2.OXO.PENTANOIC.ACID | | ND | | | | 0.25585 | |
| In-house | 130.0874 | | 1.8 | | NEG | LEUCINE.Peak.1.2 | | ND | | | | 0.04769 | Drought |
| In-house | 131.0462 | | 1.3 | | NEG | L.ASPARAGINE | | ND | | | | 0.18108 | |
| In-house | 131.0826 | | 1.2 | | NEG | L.ORNITHINE | | ND | | | | 0.20336 | |
| In-house | 132.0302 | | 1.4 | | NEG | ASPARTATE | | ND | | | | 0.60625 | |
| In-house | 134.0472 | | 1.3 | | NEG | ADENINE | | ND | | | | 0.04885 | Normal |
| In-house | 137.0244 | | 11.4 | | NEG | SALICYLATE | | ND | | | | 0.74407 | |

| | | | | | | | | | | | | | |
|---|---|---|---|---|---|---|---|---|---|---|---|---|---|
| In-house | 137.0357 | | 1.3 | | NEG | UROCANATE | | ND | | | | 0.54316 | |
| In-house | 145.0619 | | 1.3 | | NEG | L.GLUTAMINE | | ND | | | | 0.92714 | |
| In-house | 146.0459 | | 1.3 | | NEG | L.GLUTAMIC.ACID | | ND | | | | 0.01547 | Drought |
| In-house | 154.0622 | | 1.3 | | NEG | L.HISTIDINE | | ND | | | | 0.65496 | |
| In-house | 161.0456 | | 1.8 | | NEG | 3.HYDROXY.3.METHYL GLUTARATE | | ND | | | | 0.02128 | Drought |
| In-house | 163.0764 | | 14.6 | | NEG | Eugenol | | ND | | | | 0.37297 | |
| In-house | 164.0717 | | 2.2 | | NEG | L.PHENYLALANINE.Peak .1.2 | | ND | | | | 0.74209 | |
| GNPS | | 169.000 | | 1.8 | [M-H]- | | Gallate\|Pyrogallol-5-carboxylic acid\|3,4,5-Trihydroxybenzoate\|3,4,5-Trihydroxybenzoic acid\|Gallic acid | ND | | | | 0.06603 | |
| In-house | 171.0299 | | 1.5 | | NEG | 3.DEHYDROSHIKIMATE | | ND | | | | 0.08145 | |
| GNPS | | 173.046 | | 21.0 | [M-H]- | | Juarezic Acid | ND | | | | | |
| Both | 173.0819 | 173.082 | 9.8 | 9.5 | NEG | SUBERIC.ACID | RP015311 Suberic acid\|Octanedioic acid | ND | | | | 0.64230 | |
| In-house | 174.0884 | | 1.3 | | NEG | Citrulline | | ND | | | | 0.07878 | |
| GNPS | | 177.019 | | 20.1 | [M-H]- | | esculetin\|6,7-dihydroxychromen-2-one | ND | | | | 0.00988 | Drought |
| In-house | 177.0405 | | 1.4 | | NEG | D.GULONIC.ACID.GAMA .LACTONE | | ND | | | | 0.66163 | |
| In-house | 177.0921 | | 16.1 | | NEG | Methyleugenol | | ND | | | | 0.27340 | |
| In-house | 179.0561 | | 1.3 | | NEG | Sugars.Monosaccharides.Hexoses | | ND | | | | 0.12823 | |
| In-house | 181.0717 | | 1.3 | | NEG | Sugars.Alcohol.Hexoses | | ND | | | | 0.02509 | Drought |
| GNPS | | 186.114 | | 8.9 | [M-H]- | | 3-Amino-Beta-Pinene | ND | | | | | |
| Both | 187.0976 | 187.098 | 11.3 | 10.8 | NEG | AZELAIC.ACID | AZELAIC ACID | ND | | | | 0.34514 | |

| | | | | | | | | | | | | | |
|---|---|---|---|---|---|---|---|---|---|---|---|---|---|
| GNPS | | 187.134 | | 12.7 | NEG | | X10.HYDROXYDECANOATE | ND | | | | 0.83562 | |
| Both | 195.0510 | 195.051 | 1.5 | 1.4 | NEG | GLUCONIC.ACID | KNA00638 D-Gluconic acid\|D-Gluconate\|D-gluco-Hexonic acid | ND | | | | 0.56646 | |
| GNPS | | 197.009 | | 4.4 | [M-H]- | | KO001812 Syringate\|Syringic acid | ND | | | | | |
| GNPS | | 201.113 | | 12.2 | [M-H]- | | SEBACATE | 201.11322 | $C_{10}H_{18}O_4$ | 0.05 | Protein-like | 0.36816 | |
| GNPS | | 203.083 | | 3.4 | M+H | | 3 L-Tryptophan\|Tryptophan\|(S)-alpha-Amino-beta-(3-indolyl)-propionic acid | | | | | | |
| GNPS | | 215.129 | | 13.2 | [M-H]- | | Undecanedioic acid | 215.12887 | $C_{11}H_{20}O_4$ | 0.05 | Protein-like | | |
| GNPS | | 229.156 | | 2.6 | [M-H]- | | Spectral Match to Leu-Val from NIST14 | | | | | 0.05048 | |
| Both | 241.0830 | 241.083 | 2.0 | 1.8 | NEG | THYMIDINE.Peak.2 | THYMIDINE | 241.083 | $C_{10}H_{14}O_5N_2$ | -0.06 | Lignin-like | 0.93616 | |
| GNPS | | 241.145 | | 13.9 | [M-H]- | | (Z)-2-octylpent-2-enedioic acid | 241.14453 | $C_{13}H_{22}O_4$ | 0.02 | Protein-like | | |
| In-house | 242.0783 | | 1.3 | | NEG | Cytidine | | 242.078 | $C_9H_{13}O_5N_3$ | -0.001 | Lignin-like | 0.09012 | |
| GNPS | | 249.15 | | 20.4 | [M-H]- | | (4aR,5S,8aS,9aR)-9a-hydroxy-3,4a,5-trimethyl-5,6,7,8,8a,9-hexahydro-4H-benzo[f][1]benzofuran-2-one | 249.14962 | $C_{15}H_{22}O_3$ | -0.02 | Lignin-like | 0.00002 | Drought |
| In-house | 253.2173 | | 23.0 | | NEG | PALMITOLEIC.ACID | | 253.21731 | $C_{16}H_{30}O_2$ | -0.03 | Lipid-like | 0.62557 | |
| GNPS | | 265.148 | | 24.3 | [M-H]- | | Spectral Match to Dodecyl sulfate from NIST14 | | | | | | |

| | | | | | | | | | | | | | |
|---|---|---|---|---|---|---|---|---|---|---|---|---|---|
| Both | 266.0895 | 266.089 | 1.4 | 1.5 | NEG | ADENOSINE.Peak.1.2 | DEOXYGUANOSINE | 266.089 | $C_{10}H_{13}O_4N_5$ | -0.06 | Lignin-like | 0.01076 | Drought |
| In-house | 267.0735 | | 1.4 | | NEG | Inosine.Peak.1 | | 267.073 | $C_{10}H_{12}O_5N_4$ | 0.05 | Lignin-like | 0.05600 | |
| Both | 269.0455 | 269.046 | 17.4 | 17.0 | NEG | Emodin | Aloe-emodin | 269.04554 | $C_{15}H_{10}O_5$ | 0.01 | Condensed HC | 0.00001 | Drought |
| GNPS | | 271.061 | | 12.8 | [M-H]- | | 3-(2,6-dihydroxyphenyl)-4-hydroxy-6-methyl-3H-2-benzofuran-1-one | 271.06119 | $C_{15}H_{12}O_5$ | 0.03 | Lignin-like | 0.78298 | |
| GNPS | | 277.145 | | 18.7 | [M-H]- | | Mono-2-ethylhexyl phthalate from NIST14 | 277.14453 | $C_{16}H_{22}O_4$ | -0.01 | Lignin-like | | |
| GNPS | | 277.217 | | 20.5 | [M-H]- | | Pinolenic acid from NIST14 | 277.21732 | $C_{18}H_{30}O_2$ | -0.05 | Lipid-like | | |
| GNPS | | 285.041 | | 16.1 | [M-H]- | | 3',4',5,7-tetrahydroxyflavone | 285.04047 | $C_{15}H_{10}O_6$ | -0.01 | Condensed HC | | |
| GNPS | | 285.065 | | 14.8 | [M-H]- | | 3',4',5,7-tetrahydroxyflavone | | | | | | delete? |
| GNPS | | 293.176 | | 18.8 | M-H2O-H | | Pregna-4,16-diene-3,20-dione from NIST14 | 293.17582 | $C_{17}H_{26}O_4$ | 0.03 | Lipid-like | 0.13246 | |
| GNPS | | 301.217 | | 21.5 | N/A | | 5-HETE from NIST14 | 301.21730 | $C_{20}H_{30}O_2$ | 0.01 | Lipid-like | 0.15320 | |
| GNPS | | 303.233 | | 22.6 | M-H-H2O | | 5S-Hydroxy-6E,8Z,11Z-eicosatrienoic acid from NIST14 | 303.23296 | $C_{20}H_{32}O_2$ | -0.01 | Lipid-like | 0.95453 | |
| GNPS | | 305.248 | | 23.8 | [M-H]- | | cis-5,8,11-Eicosatrienoic acid from NIST14 | 305.249 | $C_{20}H_{34}O_2$ | 0.03 | Lipid-like | | |
| GNPS | | 311.169 | | 19.2 | [M-H]- | | 1-[2-methyl-6-[(2S,3R,4S,5S,6R)-3,4,5-trihydroxy-6-(hydroxymethyl)oxan-2-yl]oxyphenyl]ethanone | | | | | 0.00613 | Drought |

| | | | | | | | | | | | | |
|---|---|---|---|---|---|---|---|---|---|---|---|---|
| GNPS | | 313.035 | | 12.9 | [M-H] | | Endocrocin | 313.03537 | $C_{16}H_{10}O_7$ | 0.02 | Condensed HC | |
| GNPS | | 325.185 | | 19.6 | [M-H]- | | Dodecylbenzenesulfonic acid | | | | | |
| Both | 341.1089 | 341.109 | 1.3 | 1.3 | NEG | Sugars.Disaccharides | D-(+)-Trehalose from NIST14 | 341.10886 | $C_{12}H_{22}O_{11}$ | 0.23 | Carbohydrate | 0.22951 |
| Both | 343.0823 | 343.082 | 20.5 | 19.7 | NEG | Usnic.acid | Usnic acid | 343.08232 | $C_{18}H_{16}O_7$ | 0.01 | Lignin-like | 0.08421 |
| GNPS | | 373.093 | | 20.1 | [M-H]- | | Atranorin | 373.09289 | $C_{19}H_{18}O_8$ | -0.003 | Lignin-like | 0.30345 |
| GNPS | | 455.102 | | 1.3 | M+FA-H | | 2-(hydroxymethyl)-5-(2-oxopropyl)-7-[(2S,3R,4S,5S,6R)-3,4,5-trihydroxy-6-(hydroxymethyl)oxan-2-yl]oxychromen-4-one | | | | | |
| GNPS | | 487.343 | | 15.9 | [M-H]- | | (4aS,6aS,6bR,9R,10R,11R,12aR)-10,11-dihydroxy-9-(hydroxymethyl)-2,2,6a,6b,9,12a-hexamethyl-1,3,4,5,6,6a,7,8,8a,10,11,12,13,14b-tetradecahydropicene-4a-carboxylic acid | 487.34286 | $C_{30}H_{48}O_5$ | 0.08 | Lipid-like | |
| In-house | 503.1617 | | 1.3 | | NEG | Sugars.Trisaccharides | | 503.16174 | $C_{18}H_{32}O_{16}$ | 0.05 | Carbohydrate | 0.08278 |
| GNPS | | 547.266 | | 21.5 | M-H | | BML00943 Animicin A | 547.26598 | $C_{28}H_{40}N_2O_9$ | 0.23 | Lignin-like | |

### Table S5d. Matches to both in-house metabolomics and GNPS libraries

| **In-house m/z** | **GNPS m/z** | **In-house Library Match** | **RT (min)** | **GNPS Match** | **GNPS RT Mean (min)** | **GNPS shared fragment ions** | **DI (m/z)** | **DI Formula** | **DI Error (ppm)** |
|---|---|---|---|---|---|---|---|---|---|
| 173.0819 | 173.082 | Suberic acid | 9.84 | Massbank:RP015311 Suberic acid\|Octanedioic acid | 9.5 | 4 | ND | NA | NA |
| 187.0976 | 187.098 | Azelaic acid | 11.3 | Azelaic acid | 10.78 | 4 | ND | NA | NA |
| 195.0510 | 195.051 | D-Gluconic acid | 1.52 | Massbank:KNA00638 D-Gluconic acid\|D-Gluconate\|D-gluco-Hexonic acid | 1.43 | 6 | ND | NA | NA |
| 241.0830 | 241.083 | Thymidine | 1.96 | Thymidine | 1.85 | 4 | 241.08301 | $C_{10}H_{14}O_5N_2$ | -0.06 |
| 266.0895 | 266.089 | Adenosine | 1.38 | Deoxyguanosine | 1.46 | 4 | 266.08949 | $C_{10}H_{13}O_4N_5$ | -0.06 |
| 269.0455 | 269.046 | Emodin | 17.41 | Massbank:LU108653 Aloe-emodin | 16.98 | 3 | 269.04600 | $C_{15}H_{10}O_5$ | 0.01 |
| 341.1089 | 341.109 | Disaccharides/Trehalose | 1.345 | Spectral Match to D-(+)-Trehalose from NIST14 | 1.32 | 7 | 341.10900 | $C_{12}H_{22}O_{11}$ | 0.23 |
| 343.0823 | 343.082 | Usnic acid | 20.46 | Massbank:BML00262 Usnic acid | 19.71 | 4 | 343.08200 | $C_{18}H_{16}O_7$ | 0.01 |

### Table S6. Precursor masses in polyphenol subnetwork (Figure 5b) matched to GNPS or detected with DI-FTICR-MS

| GNPS Precursor mass | GNPS Adduct | GNPS Library Match | Shared Peaks, Score | GNPS Error (ppm) | DI *m/z* | *DI Extract | DI formula (N<=2, S,P = 0) | DI Error (ppm) | Element al Ratio Class | MS2 only in | Drought> Control P<0.05 |
|---|---|---|---|---|---|---|---|---|---|---|---|
| 177.019 | [M-H]- | Massbank: esculetin\|6,7-dihydroxychromen-2-one | 3, 0.92 | 0.3954 | <200 *m/z* | ND | NA | NA | ConHC | Drought | 9.88E-3 |
| 197.009 | [M-H]- | Massbank:KO001812 Syringate\|Syringic acid | 3 ,0.80 | -45.685 | <200 *m/z* | ND | NA | NA | Lignin | Drought | |
| 209.155 | | | | | 209.15471 | All | C13H22O2 | -0.009 | Lipid | | 3.52E-3 |
| 211.046 | | | | | 211.04634 | MeOH | NA | NA | NA | | 3.96E-6 |
| 225.008 | | | | | 225.00777 | All | NA | NA | NA | | |
| 225.150 | | | | | 225.14963 | All | C13H22O3 | -0.032 | Lipid | Drought | |
| 227.071 | | | | | 227.07139 | All | C14H12O3 | -0.102 | Lignin | | |
| 239.060 | | | | | 239.05965 | All | NA | NA | NA | | |
| 239.129 | | | | | 239.12889 | All | C13H20O4 | -0.015 | Protein | | 7.85E-5 |
| 241.145 | [M-H]- | (Z)-2-octylpent-2-enedioic acid | 3, 0.78 | -3.318 | 241.14453 | MeOH, $CHCl_3$ | C13H22O4 | 0.078 | Protein | | |
| 249.040 | | | | | 249.04046 | All | C12H10O6 | 0.003 | Lignin | Drought | |
| 249.150 | [M-H]- | (4aR,5S,8aS,9aR)-9a-hydroxy-3,4a,5-trimethyl-5,6,7,8,8a,9-hexahydro-4H-benzo[f][1]benzofuran-2-one | 5, 0.89 | -0.803 | 249.14961 | All | C15H22O3 | 0.028 | Lignin | Drought | 2.14E-5 |
| 253.144 | | | | | 253.14386 | All | NA | NA | NA | | 4.41E-3 |
| 253.217 | | In-house:Palmitoleic Acid | N/A | N/A | 253.21731 | All | C16H30O2 | -0.028 | Lipid | | |
| 255.160 | | | | | 255.16018 | $H_2O$, MeOH | C14H24O4 | 0.013 | Lipid | | |
| 261.040 | | | | | 261.04047 | $H_2O$, MeOH | C13H10O6 | -0.018 | ConHC | Drought | |
| 263.056 | | | | | 263.05613 | $H_2O$, MeOH | C13H12O6 | -0.055 | Lignin | Drought | 1.23E-2 |
| 267.233 | | | | | 267.23297 | $H_2O$, MeOH | C17H32O2 | -0.061 | Lipid | | |

| | | | | | | | | | | | |
|---|---|---|---|---|---|---|---|---|---|---|---|
| 269.046 | [M-H]- | Massbank:LU108653 Aloe-emodin | 3, 0.79 | 0.283 | 269.04553 | All | C15H10O5 | 0.066 | ConHC | | 9.01E-6 |
| 277.004 | | | | | 277.00400 | All | N/A | NA | | Drought | 1.62E-2 |
| 279.233 | | | | | 279.23294 | MeOH, $CHCl_{l3}$ | C18H32O2 | 0.033 | Lipid | | |
| 281.249 | | | | | 281.24860 | All | C18H34O2 | 0.012 | Lipid | | |
| 283.001 | | | | | 283.00149 | MeOH | N/A | | | Drought | |
| 283.192 | | | | | 283.19151 | $H_2O$, MeOH | C16H28O4 | -0.084 | Lipid | | |
| 285.041 | [M-H]- | 3',4',5,7-tetrahydroxyflavone | 6, 0.80 | 14.033 | 285.04047 | All | C15H10O6 | -0.038 | ConHC | | |
| 291.066 | | | | | 291.06614 | H2O | C18H12O4 | 0.490 | ConHC | Drought | 4.35E-2 |
| 291.124 | | | | | 291.12378 | All | C16H20O5 | 0.057 | Lignin | Drought | |
| 301.072 | | | | | 301.07173 | All | C16H14O6 | 0.094 | Lignin | | |
| 302.985 | | | | | 302.98497 | MeOH | NA | | | | 2.44E-5 |
| 305.248 | [M-H]- | Spectral Match to cis-5,8,11-Eicosatrienoic acid from NIST14 | 5, 0.80 | 3.276 | 305.24860 | $CHCl_{l3}$ | C20H34O2 | 0.027 | Lipid | Drought | |
| 313.035 | [M-H]- | Endocrocin | 3, 0.84 | 5.111 | 313.03536 | $H_2O$, MeOH | C16H10O7 | 0.060 | ConHC | Drought | |
| 315.015 | | | | | 315.01463 | $H_2O$, MeOH | C15H8O8 | 0.034 | ConHC | Drought | |
| 329.103 | | | | | 329.10306 | All | C18H18O6 | 0.017 | Lignin | Drought | 2.82E-2 |
| 353.142 | | | | | 353.14240 | MeOH | NA | | | | |
| 363.036 | | | | | 363.03572 | All | C16H12O10 | 0.136 | ConHC | Drought | |
| 367.082 | | | | | 367.08230 | All | C20H16O7 | 0.082 | Lignin | Drought | |
| 389.254 | | | | | 389.25442 | $H_2O$, MeOH | C20H38O7 | 0.151 | Protein | | 1.18E-4 Control> Drought |
| 403.046 | | | | | 403.04578 | All | C22H12O8 | 0.407 | ConHC | | |
| 403.082 | | | | | 403.08230 | MeOH, $CHCl_{l3}$ | C23H16O7 | 0.064 | ConHC | | 2.77E-2 Control> Drought |
| 405.062 | | | | | 405.06153 | All | C22H14O8 | 0.145 | ConHC | | |
| 437.052 | | | | | 437.05166 | $H_2O$, | NA | NA | | Drought | |
| 453.046 | | | | | 453.04619 | All | C22H14O11 | 0.310 | ConHC | | |
| 469.078 | | | | | 469.07759 | All | C23H18O11 | 0.089 | ConHC | | |
| 471.312 | | | | | 471.31152 | MeOH, $CHCl_{l3}$ | C29H44O5 | 0.163 | Lipid | Drought | |

* All = detected in all three (water, methanol and chloroform) extracts.

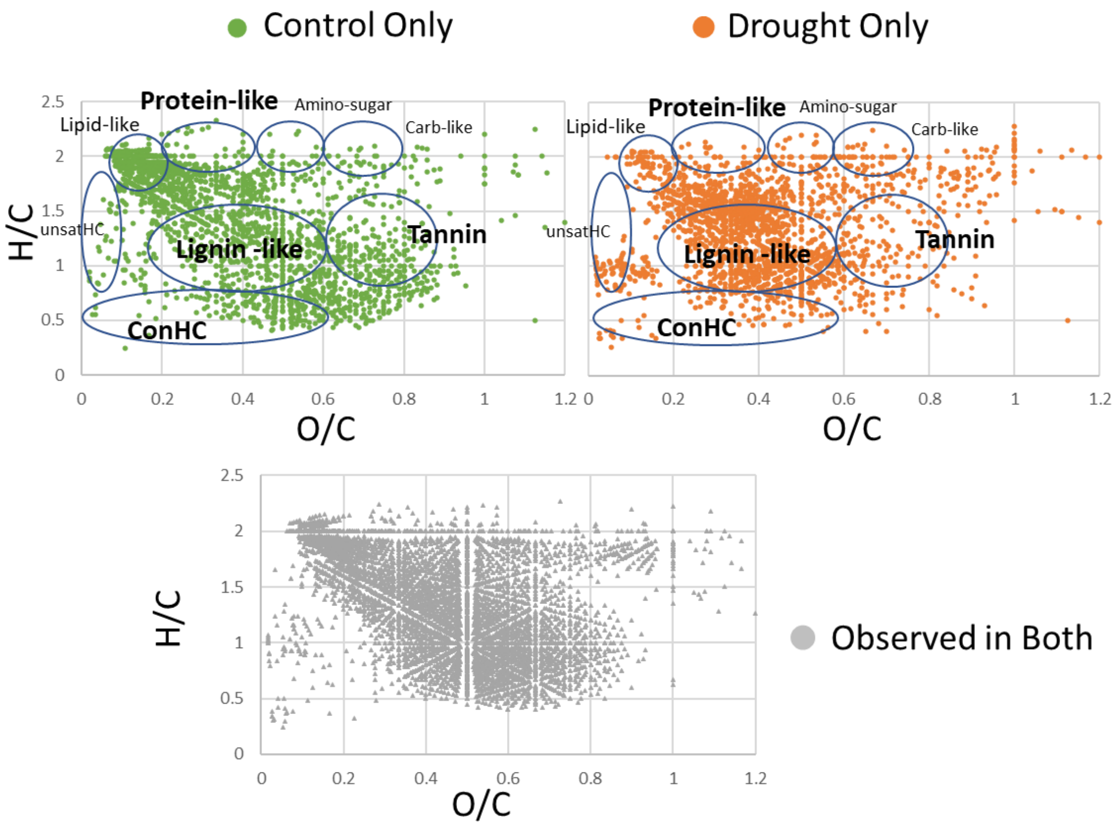

Figure S1. van Krevelen diagrams for drought and control detritusphere SOM at week 12

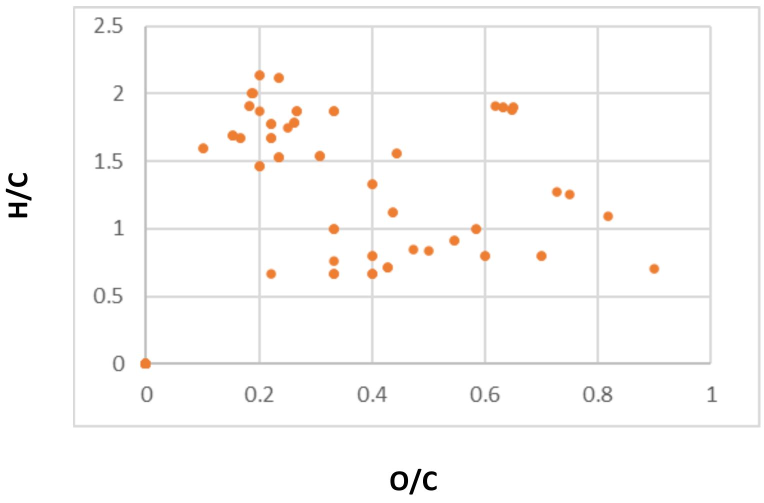

Figure S2. van Krevelen plot of FTICR-MS molecular formulas also detected with LC-MS at significantly higher abundance in the drought (primarily CHO)

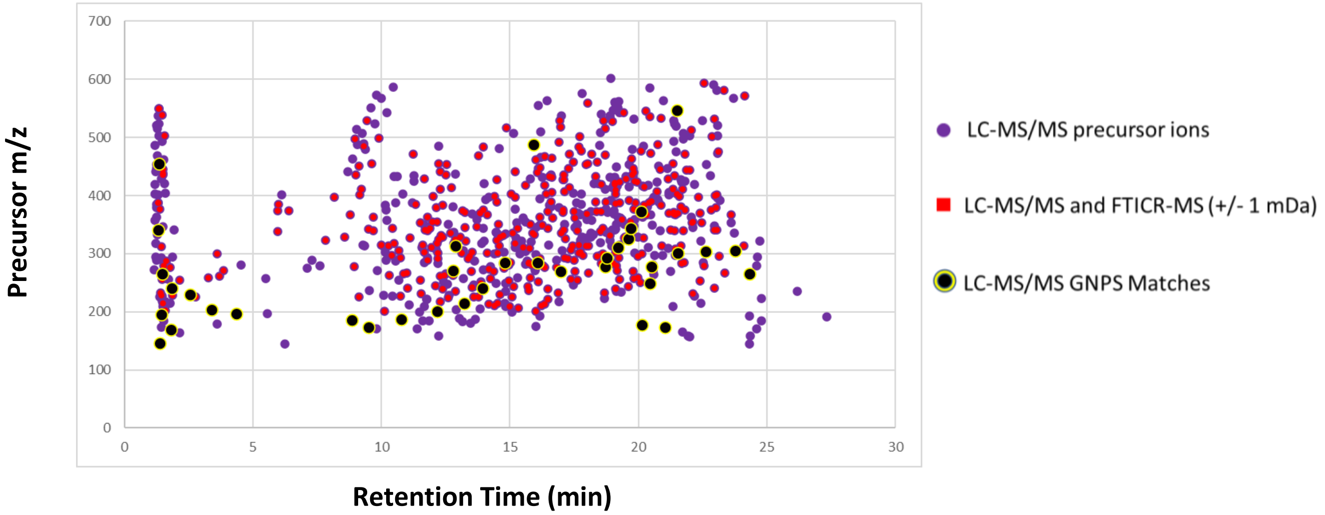

Figure S3. LC-MS/MS chromatogram of precursor retention times vs *m/z* for which fragment ion spectra were acquired in negative ion mode at week 12. This is overlayed with the same masses (+/- 1 mDa) detected with DI-FTICRMS. Precursor masses with negative mode fragment ion spectra that matched to GNPS library spectra with a similarity score of 0.7 or greater are highlighted in black and listed in Table S4c.

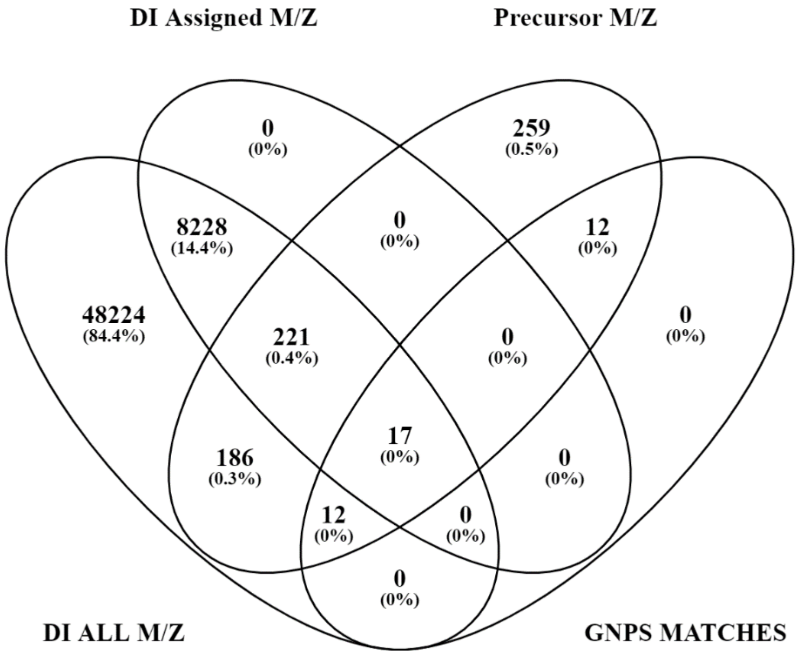

Figure S4. Venn Diagram of DI-FTICRMS *m/z* detected as precursor masses and assigned molecular formula or matched to GNPS libraries.

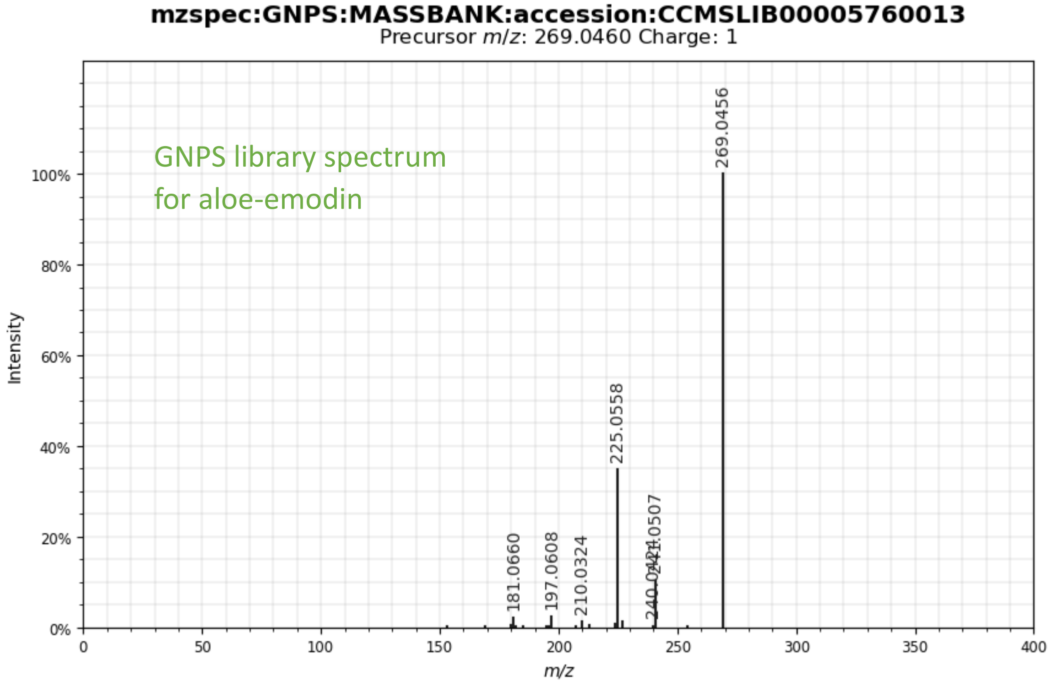

a)

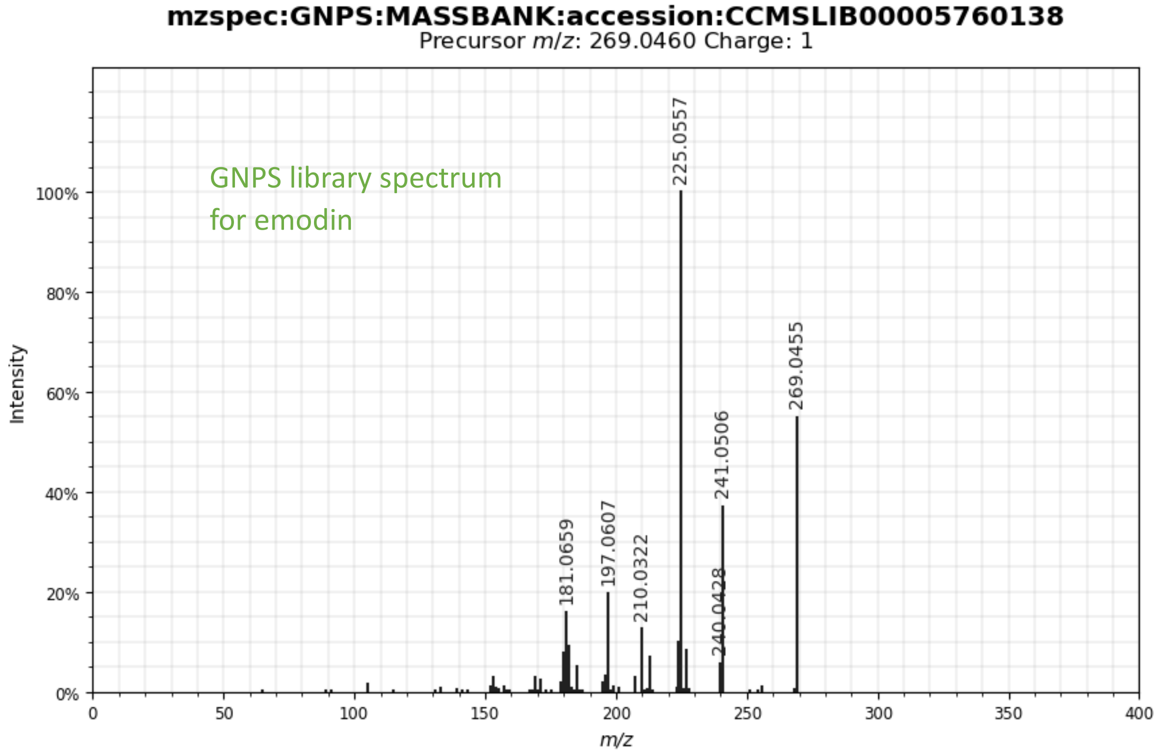
mzspec:GNPS:MASSBANK:accession:CCMSLIB00005760138
Precursor m/z: 269.0460 Charge: 1
GNPS library spectrum
for emodin
Intensity
m/z
225.0557
269.0455
241.0506
240.0428
210.0322
197.0607
181.0659

b)

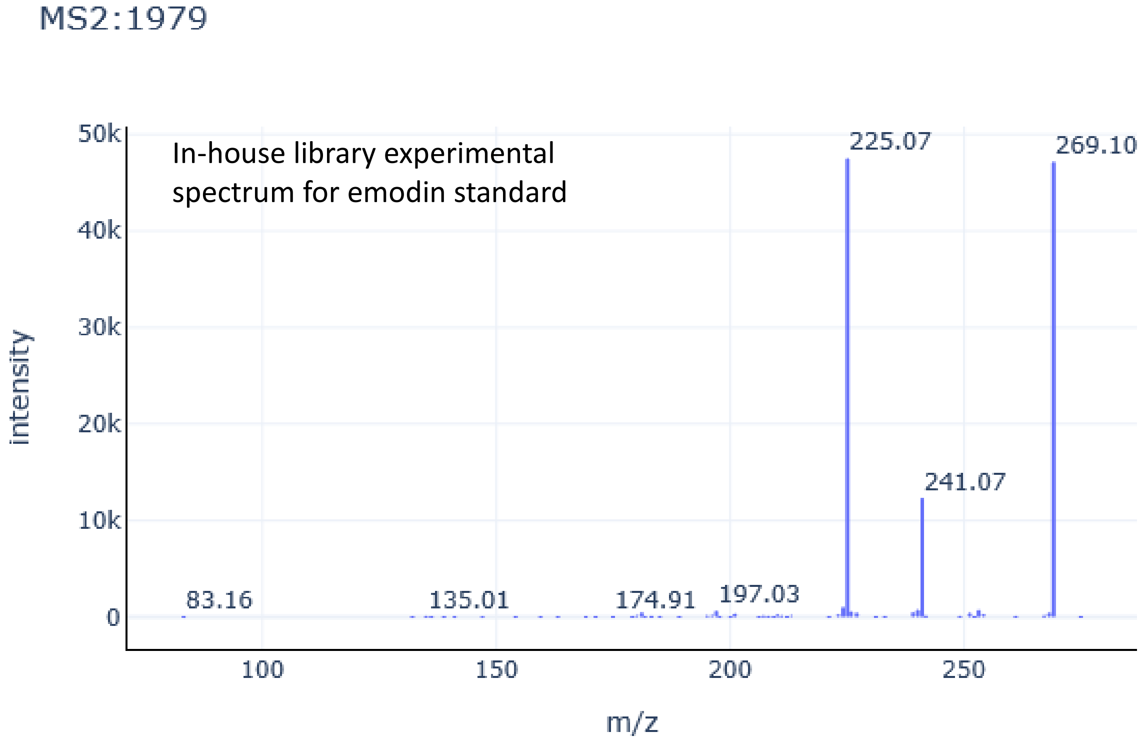
MS2:1979
In-house library experimental
spectrum for emodin standard
intensity
m/z
225.07
269.10
241.07
83.16
135.01
174.91
197.03

c)

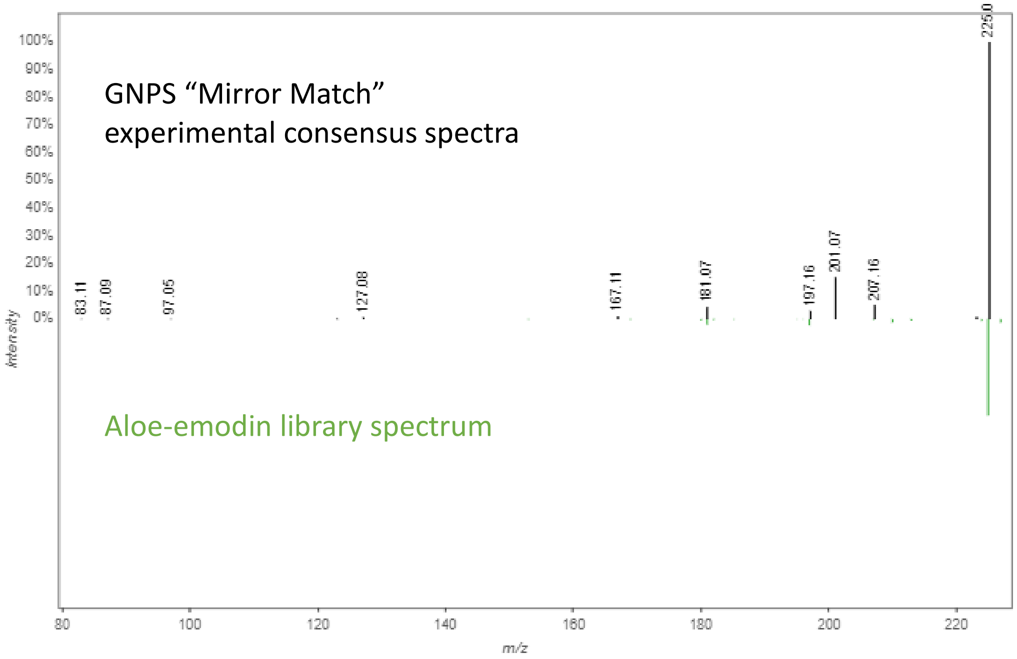

d)

Figure S5a-d. Negative mode GNPS library fragmentation spectra for a) aloe-emodin and b) emodin, c) in-house library experimental spectrum of emodin standard, and d) mirror match spectra of averaged experimental consensus spectra to GNPS library match spectra for aloe-emodin. Experimental tandem mass spectra used to generate the composite consensus spectra for comparison with library spectra included 12 experimental spectra (scans 1250-1810) collected between 16.95 and 17.02 minutes from 6 samples (2 replicates each from control and 2 from drought) in which m/z 269.046 was detected and fragmented. Emodin standard was detected at 17.41 minutes using the in-house library and processing method.

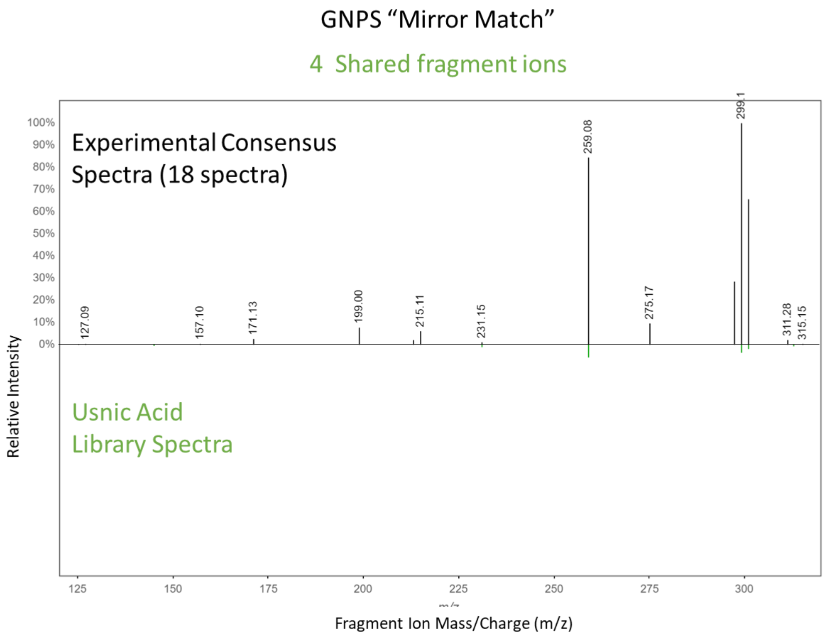

e)

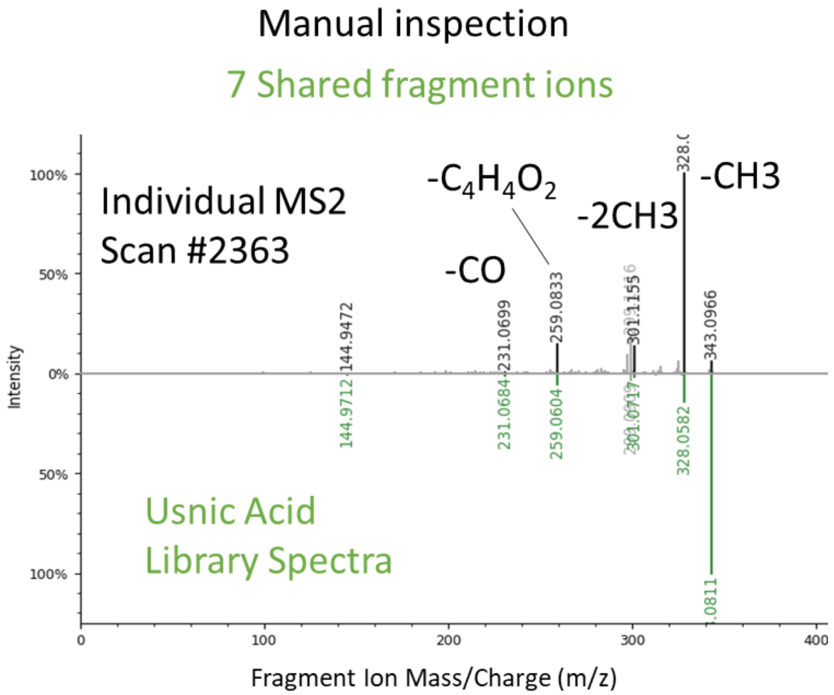

f)

Figure S5e-f. GNPS mirror match of e) usnic acid library spectra and experimental consensus spectra generated by GNPS; f) manual mirror match between individual scan 2362 and usnic acid library spectra.

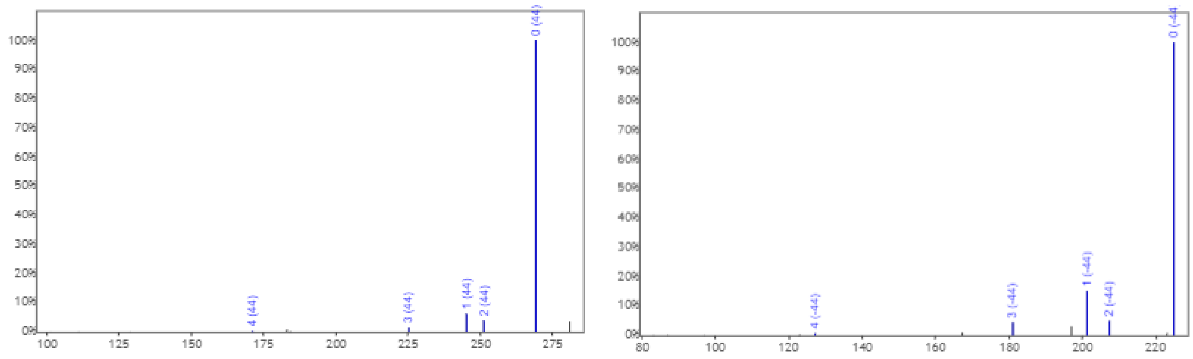

Figure S5g. Aligned fragment ion spectra for endocrocin (FTICRMS ConHC)(left) and aloe-emodin (right)

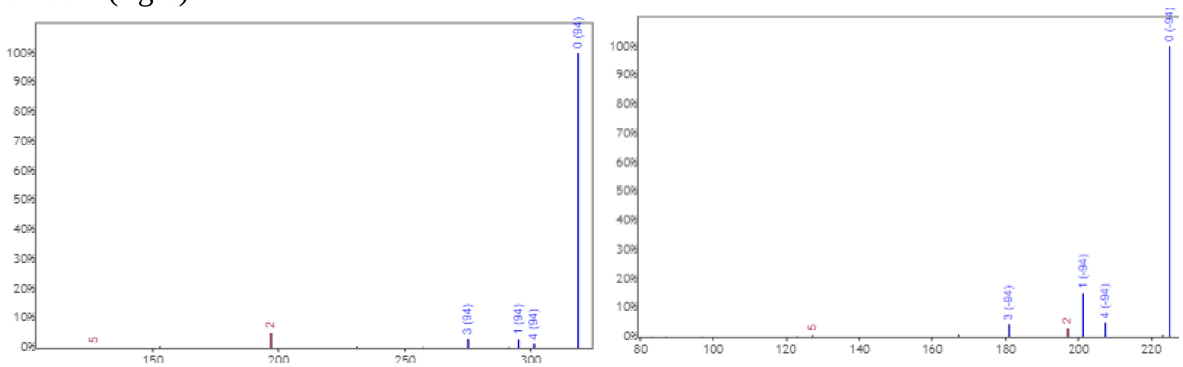

Figure S5h. Aligned fragment ion spectra for *m/z* 363.036 (FTICRMS ConHC) (left) and aloe-emodin (right) (fragment *m/z* 127 and 197 not shifted)

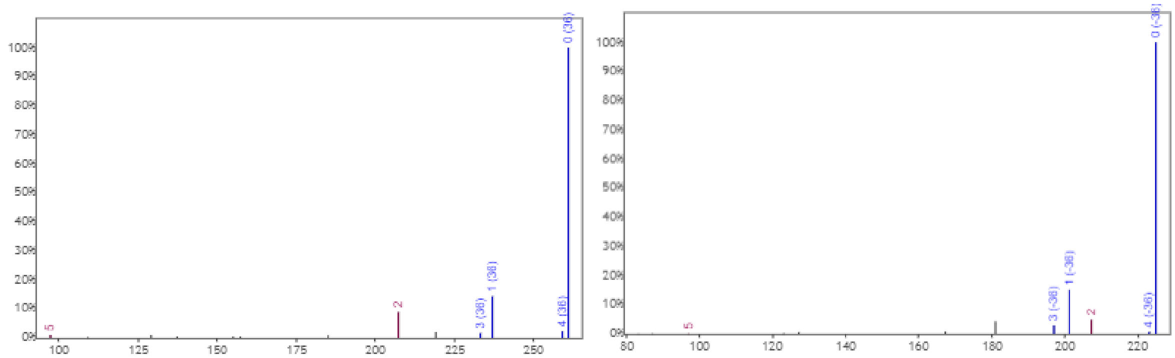

Figure S5i. Aligned fragment ion spectra for *m/z* cis-5,8,11-Eicosatrienoic acid (FTICRMS assigned lipid-like)(left) and aloe-emodin (right) (fragment *m/z* 97 and 207 not shifted)

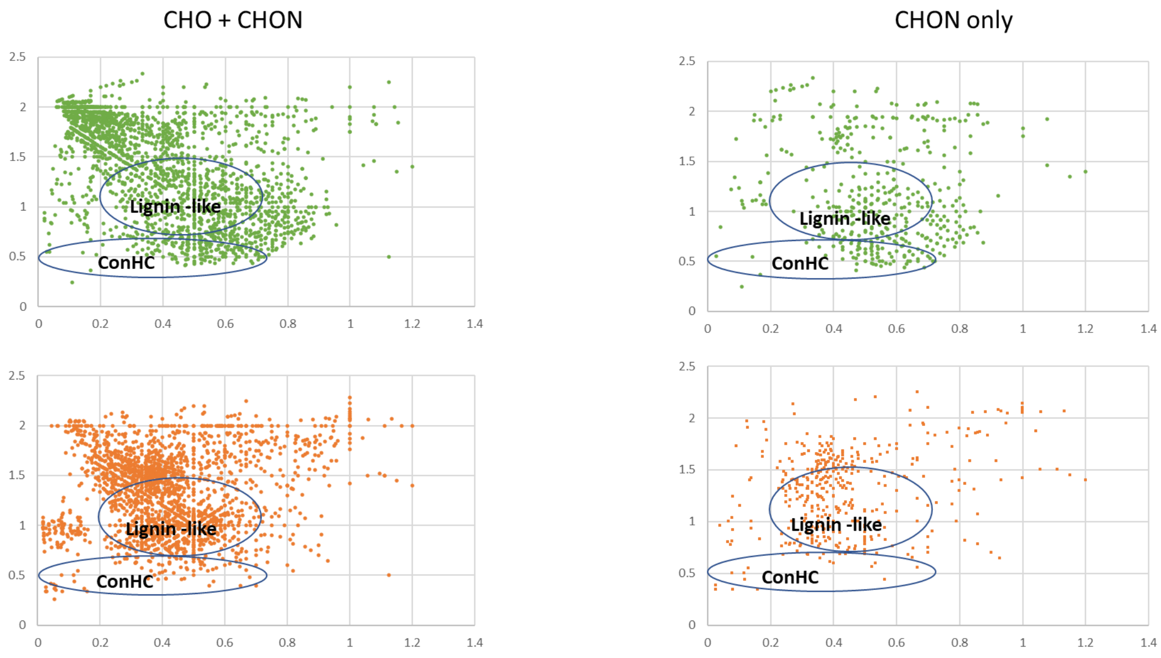

Figure S6. Molecular formulas detected with DI-FT-ICR-MS at week 12 unique to drought (bottom, orange) and control (top, green), CHO and CHON (left) and CHON only (right)

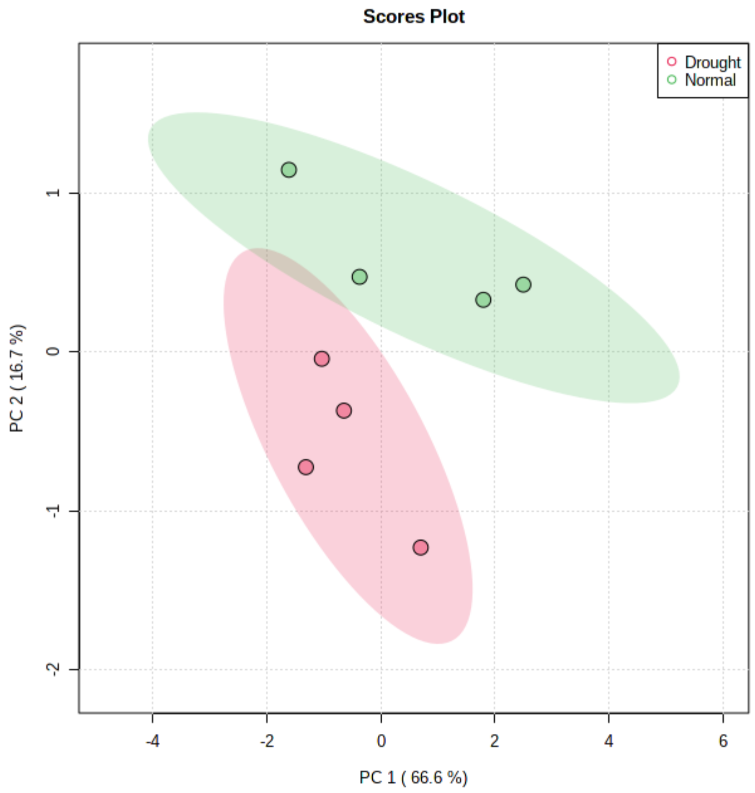

a)

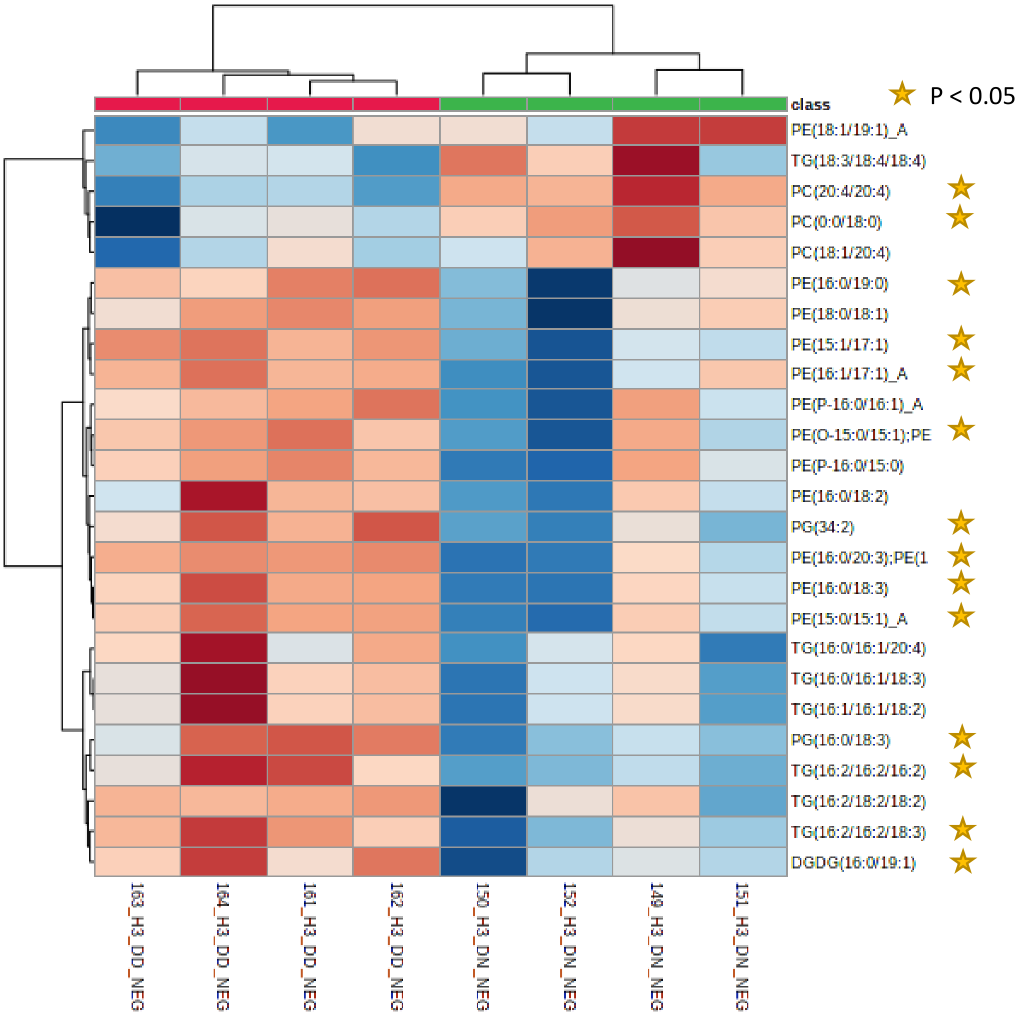

b)

Figure S7. a) PCA plot of lipids identified in negative mode from detritusphere soil, drought and control at 12 weeks (Harvest 3). b) Heatmap of Top 25 most significant lipids ($P<0.08$) from negative and positive ion mode. Lipids with the most significantly different abundances ($P<0.05$) in drought and control are starred.
